# Identifying Behavioral Structure from Deep Variational Embeddings of Animal Motion

**DOI:** 10.1101/2020.05.14.095430

**Authors:** Kevin Luxem, Petra Mocellin, Falko Fuhrmann, Johannes Kürsch, Stefan Remy, Pavol Bauer

## Abstract

Quantification and detection of the hierarchical organization of behavior is a major challenge in neuroscience. Recent advances in markerless pose estimation enable the visualization of highdimensional spatiotemporal behavioral dynamics of animal motion. However, robust and reliable technical approaches are needed to uncover underlying structure in these data and to segment behavior into discrete hierarchically organized motifs. Here, we present an unsupervised probabilistic deep learning framework that identifies behavioral structure from deep variational embeddings of animal motion (VAME). By using a mouse model of beta amyloidosis as a use case, we show that VAME not only identifies discrete behavioral motifs, but also captures a hierarchical representation of the motif’s usage. The approach allows for the grouping of motifs into communities and the detection of differences in community-specific motif usage of individual mouse cohorts that were undetectable by human visual observation. Thus, we present a novel and robust approach for quantification of animal motion that is applicable to a wide range of experimental setups, models and conditions without requiring supervised or a-priori human interference.

## 1 Introduction

The brain is a dynamical system and its dynamics are reflected in the actions it performs. Thus, observable motion is a valuable resource for understanding brain function. In most of the current neuroethological studies this resource has only been partially utilized (Krakauer, Ghazanfar, Gomez-Marin, MacIver, & Poeppel, 2017). Reaching the goal of maximizing information content requires a complete capture of observable motion and unbiased interpretation of behavioral complexity. Unsupervised methods provide a gateway for this purpose as they do not rely on human annotations like their counterparts, supervised methods (Nilsson et al., 2020; Segalin et al., 2021; Bohnslav et al., 2021). Moreover, unsupervised methods are able to learn rich dynamical representations of behavior on a sub-second scale, which are otherwise not detectable (Gomez-Marin, Paton, Kampff, Costa, & Mainen, 2014; Anderson & Perona, 2014; Wiltschko et al., 2015; Berman, 2018; Brown & de Bivort, 2018; Datta, Anderson, Branson, Perona, & Leifer, 2019). The need for unsupervised behavioral quantification methods has been recently recognized and innovative approaches in this direction have been introduced (Wiltschko et al., 2015; Berman, Choi, Bialek, & Shaevitz, 2014; Hsu & Yttri, 2021). While there is a broad agreement among researchers in computational ethology that observable behavior can be encoded in a lower dimensional subspace or manifold (Wiltschko et al., 2015; Brown & de Bivort, 2018; Berman, 2018; Datta et al., 2019), current methods insufficiently capture the complete spatiotemporal dynamics of behavior (Datta et al., 2019).

Recently, pose estimation tools such as *DeepLabCut* (Mathis et al., 2018), *LEAP* (T. D. Pereira et al., 2019) and *DeepPoseKit* (Graving et al., 2019) enabled efficient tracking of animal body-parts via supervised deep learning. The robustness of deep neural networks allows for a high degree of generalization between datasets (Mathis et al., 2018). However, while such tools provide a continuous representation of the animal body motion, the extraction of underlying discrete states as a basis for quantification remains a key challenge.

To address this challenge and provide a reliable and robust solution, we here developed Variational Animal Motion Embedding (VAME), an unsupervised probabilistic deep learning framework for discovery of underlying latent states in behavioral signals obtained from pose estimation tools or dimensionality reduced video information (Shi et al., 2021). The input signal is learned and embedded into a lower dimensional space via a variational recurrent neural network autoencoder. Given the low dimensional representation, a Hidden-Markov-Model (HMM) learns to infer hidden states, which represent behavioral motifs. A major advantage of VAME is the ability to learn a disentangled representation of latent factors via the variational autoencoder (VAE) framework (Kingma & Welling, 2014; Rezende, Mohamed, & Wierstra, 2014; Bengio, Courville, & Vincent, 2013). This allows the model to embed a complex data distribution into a simpler prior distribution, where segments of the behavioral signal are grouped by their spatiotemporal similarity. We use a powerful autoregressive encoder to disentangle latent factors of the input data. Our approach is inspired by recent advances in the field of temporal action segmentation (Kuehne, Richard, & Gall, 2020), representation learning (Chung et al., 2015; Chen et al., 2016; Higgins et al., 2017; Jiang, Zheng, Tan, Tang, & Zhou, 2017) and unsupervised learning of multivariate time series (J. Pereira & Silveira, 2019; Ma, Zheng, Li, & Cottrell, 2019).

In this manuscript, we introduce the VAME model and workflow based on behavioral data obtained from a bottom-up recording system in an open-field. We demonstrate the capability of VAME to identify the motif structure, the hierarchical connection of motifs and their transitions in a use case of Alzheimer transgenic mice (Jankowsky et al., 2004). Within this use case, VAME is capable of detecting differences between the transgenic and control group, while no differences were detectable by human observers. In addition, we compare VAME to current state-of-the-art methods (Berman et al., 2014; Wiltschko et al., 2015) on a benchmark dataset.

## 2 Results

### 2.1 VAME: Variational Animal Motion Embedding

In our experimental setup for investigation of behavioral structures, we let mice move freely in an open-field arena (Figure 1 A). During the experiment the movement was continuously monitored with a bottom-up camera for 25 minutes (*N* = 90000 frames). In order to identify the postural dynamics of the animal from the video recordings we used DeepLabCut (DLC) (Mathis et al., 2018), a markerless pose estimation tool. For tracking, we used six virtual markers, which were positioned on all four paws of the animal, the nose, and the tailbase (Figure 1 A). We aligned the animal from its allocentric arena coordinates to its egocentric coordinates. For this, each frame was rotated around the center between nose and tail, so that the animal was aligned from left (tailbase) to right (nose) (Figure 1 A). This resulted in a time-dependent series **X** ∈ ℝ^*N×m*^ with *m* =10 (x, y)-marker positions that captured the kinematic of specified body parts. Our aim was to extract useful information from the time series data, allowing for an effective behavioral quantification given spatial and temporal information of body dynamics. Trajectory samples **x**_*i*_ ∈ ℝ^*m×w*^ (with *w* = 30) were pre defined time windows that were randomly sampled from **X** and represented the input that was used to train the VAME model. Our first goal was to identify behavioral motifs, which we defined according to (Datta et al., 2019) as “stereotyped and re-used units of movements”. The second goal was the identification of the hierarchical and transition structure of motifs aiming at the detection of patterns within these transitions.

**Figure 1.**
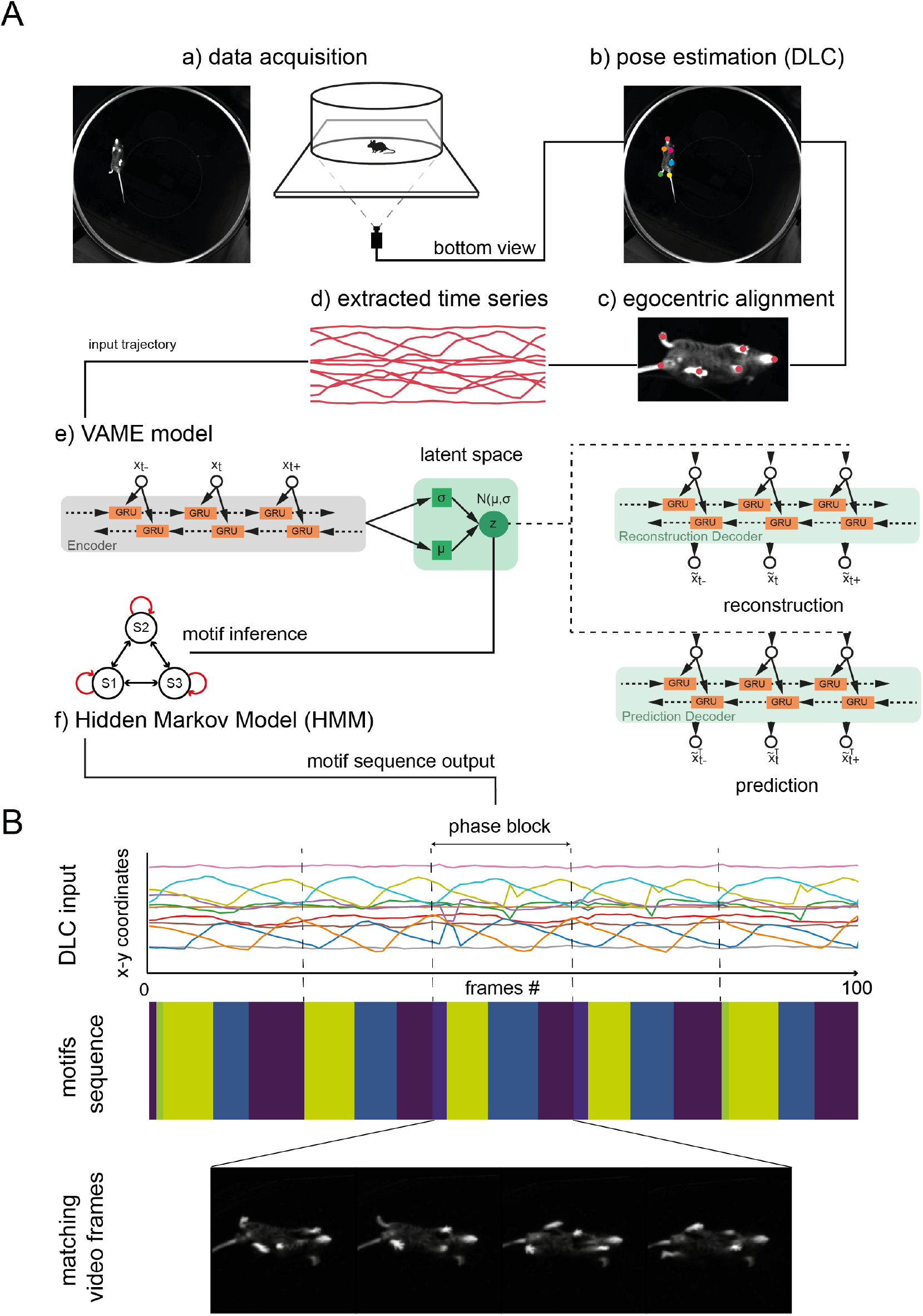
VAME: a deep learning model for unsupervised spatiotemporal behavior quantification. (A) VAME workflow. Data acquisition via bottom-up camera setup for precise body and limb kinematics. Pose estimation of bottom view (DLC). Frames are egocentrically aligned and trajectory samples are fed into the recurrent neural network model. The fully trained model resembles a dynamical system from which motifs are inferred via a Hidden-Markov-Model. (B) Top: Example trace of an egocentric aligned DLC sequence showing a full walking cycle (phase block). Middle: Corresponding motif sequence identified by VAME where the phase block structure is visible. Bottom: Matching video frames from the walking cycle.

The VAME model consists of three bidirectional recurrent neural networks (biRNN) (Schuster & Paliwal, 1997) with gated recurrent units (GRUs) (Cho et al., 2014) as basic building blocks. In our model, the encoder biRNN receives a trajectory sample **x**_*i*_ (i.e. 500 ms of behavior) and embeds the relevant information into a lower dimensional latent space **Z**. Learning is achieved by passing **z**_*i*_ ∈ ℝ^*d*^ (*d < m × w*) to a biRNN decoder which decodes the lower dimensional vector into an approximation 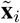 of the input trajectory. Additionally, another biRNN decoder learns to anticipate the structure of the subsequent time series trajectory 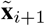 from **z**_*i*_, thereby regularizing **Z** and forcing the encoder to learn a richer representation of the behavior (Srivastava, Mansimov, & Salakhudinov, 2015). The model is trained as variational autoencoder (Kingma & Welling, 2014) with a standard normal prior (see Methods 4.3 for details). Within the VAE framework, it is possible to investigate if the model has learned a useful representation of the input data by drawing random samples from the latent space and comparing them to a subset of reconstructions (see supplemental material section 5.8). After the model is trained on the experimental data (1.3 × 10^6^ data points), the encoder embeds the data during inference onto a learned latent space. We then segment the continuous latent space into discrete behavioral motifs using a Hidden-Markov-Model (HMM) (Rabiner & Juang, 1986), thereby treating the underlying dynamical system as a discrete-state continuous-time Markov chain (Methods 4.4). See supplemental section 5.4 for more details on model selection, where we also compare the HMM against a k-means algorithm for cluster detection.

Figure 1 (B, top) depicts an example of an egocentrically aligned DLC time series (100 data points). Here, we defined a phase block, which is characterized by a full walking cycle (e.g. orange line). As shown in Figure 1 (B, middle) the inferred motif sequence aligns to the walking pattern, where the onset of each motif matches to a particular phase of the input signal. The video frames (Figure 1 (B, bottom)) display the corresponding walk cycle within the phase block.

### 2.2 Identification of behavioral motif structure

To identify motif structure and to demonstrate the power of our approach, we used four transgenic (tg) mice displaying beta-amyloidosis in cortical and hippocampal regions. These mice harbor human mutations in the APP and presenilin 1 gene (APP/PS1) (Jankowsky et al., 2004). We compared these mice to four wildtype (wt) mice housed in identical conditions. For these APP/PS1 mouse line, several behavioral differences have been reported (Huang et al., 2016). Among them, motor and coordination deficits (Onos et al., 2019), changes in anxiety levels (Lalonde, Kim, & Fukuchi, 2004) and spatial reference memory deficits (Janus, Flores, Xu, & Borchelt, 2015) were most prominently observed. This dataset forms an ideal use-case for the purpose of unsupervised behavior quantification as specific (repetitive) behavioral task settings were required to detect these differences and no differences could be detected in open field tests (Giovannetti et al., 2018).

We determined general locomotor dependent variables to investigate whether the animals show explicit locomotor differences, in particular we investigated speed, travelled distance and time spent in the center (Figure 2 A). The average speed during the trial was 6.12 ± 1.36 cm/s for wt animals and 6.84 ± 1.57 cm/s for tg animals with a maximum velocity of 50.61 ± 12.47 cm/s for wt and 57.14 ± 8.91 cm/s for tg animals. The average time spent in the center (calculated from center crossings) is 9.92 ± 1.81 seconds for wt and 17.14 ± 7.79 seconds for tg animals. Lastly, the average distance travelled was 9187.44 ± 1266.4 cm and 9937.07 ± 1367.08 cm for tg and wt respectively. No statistically significant differences were found between the groups for all measures. Nevertheless, we see a tendency for the tg group to move at a higher speed and spend more time in the center as already shown by others (Webster, Bachstetter, & Van Eldik, 2013; Biallosterski et al., 2015; Giovannetti et al., 2018)

To specifically find out whether behavioral differences were detectable by human observation in our experimental setting, we performed human classification, which was carried out by eleven human experts, all trained in behavioral neuroethology. This classification was based exclusively on the video recordings that were used for our VAME approach. We constructed an online questionnaire for blinded classification where each participant was allowed to watch all videos for an unlimited amount of time before making their decision (see Methods 4.6). Experts with previous experience about APP/PS-1 had a slightly higher classification accuracy (50.98% ± 11.04% for experts, 42.5% ± 15.61% for non-experts). The overall classification accuracy was at chance level for all participants (46.61% ± 8.41%, Figure 2 B). A similar level of behavioral homogeneity between the two animal groups was reported previously (Webster et al., 2013).

To identify behavioral structure, we applied VAME to the entire cohort of animals and inferred the latent representation for each animal. The latent dimension size was set empirically to **z** ∈ ℝ^30^, while comparing the difference between input and reconstructed signals. This parameter controls the amount of information flow between the encoder and decoder networks. By keeping this bottleneck small, the encoder learns to extract salient features from the input signal. We summarize important choices of recording and hyperparameter settings in Table S.3. Using a HMM on the latent representation, we inferred 50 motifs per animal (see section 5.5 for details) and created a hierarchical tree representation 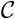 (Figure 2, C) (see Methods 4.4). By comparing branches of the tree with corresponding motif videos, we identified communities of similar behavioral motifs. We found nine behavioral communities denoted from *a* to *i*. Each community represents a cluster of movements that can be simplified into actions like e.g. rearing, turning, walking. Motifs within each community can be interpreted as a subset 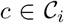 (with *i* = *a*,..., *i*) of these actions. Therefore, communities detected by VAME display a multiscale behavioral representation. All communities were visualized with their respective DLC trace and further described in the supplemental materials section 5.7. Furthermore, we provide supplemental videos (6-14) for all communities.

**Figure 2:**
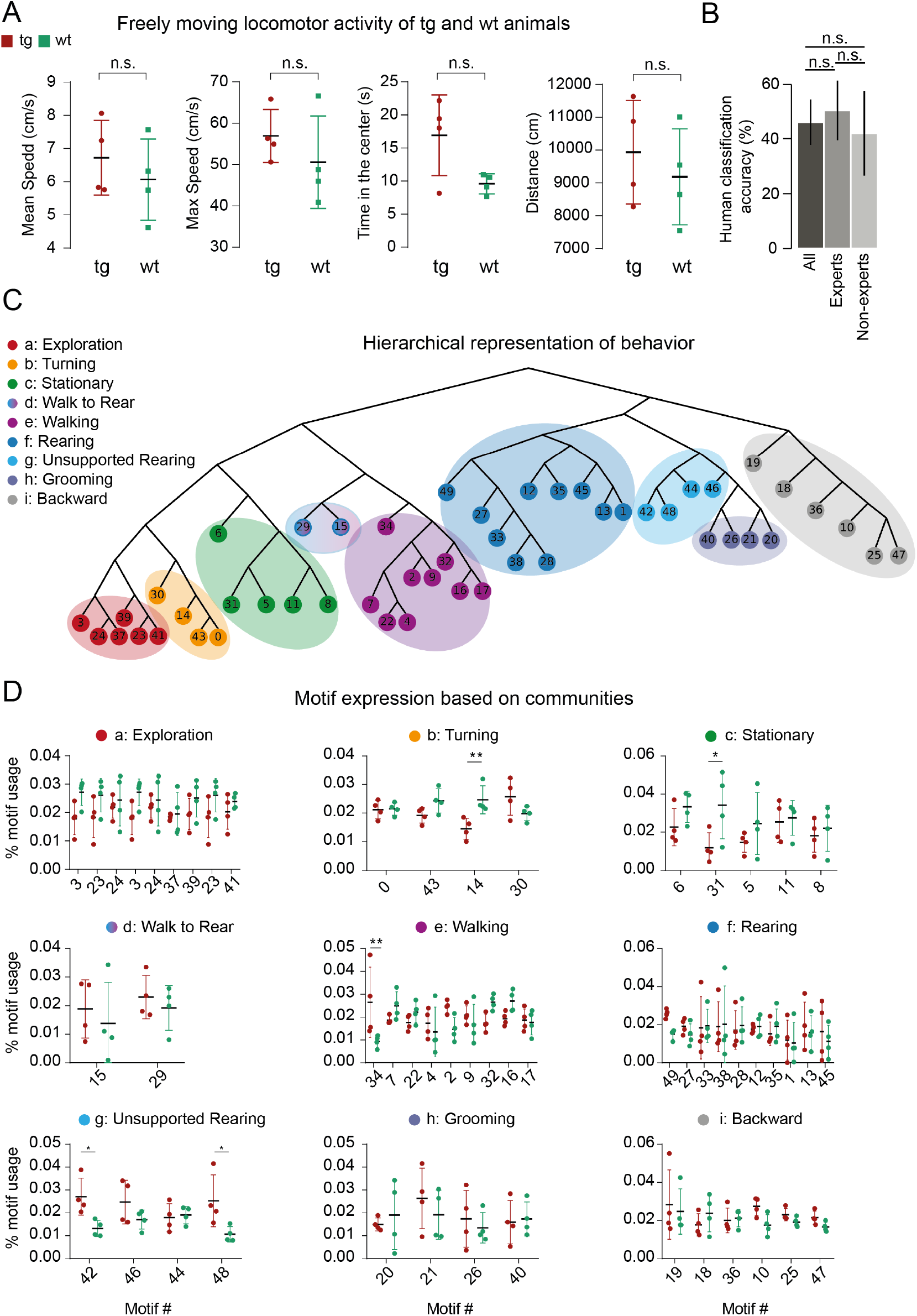
Behavioral quantification with VAME and hierarchical community clustering. (A) Locomotor activity of tg (N=4) and wt (N=4) animals. (B) Human classification accuracy. Experts refers to previous experience with APP/PS-1 animals. (C) Hierarchical representation of behavioral motifs. Color grouping on the tree are depicting communities of motifs which belong to the same observable category of behavior. (D) Quantification of motif usage ordered by communities. Differences between the tg and wt phenotype are in community b, c, e and g. (D) Motif usage between wt and tg animals in percent (%) for each community.

Next, we were interested if our model can identify differences between the APP/PS1 tg and wt mice. We compared motifs at the community level between both groups (Figure 2 D). A multiple t-Test was used and statistical significance was determined using the Holm-Sidak method with *alpha* = 0.05 (* = *P* ≤ 0.05, ** = *P* ≤ 0.01). We found a significant differences for five motifs (Table 1). This showed that the *Turning* and *Stationary* motifs increased in wt animals while tg animals showed a higher expression of *Unsupported Rearing* and *Walking* motifs, displayed by arrows in the Table 1. These motifs are described in supplemental Section 5.1 and displayed in the supplemental videos 1-5 as well as in supplemental Figure S.1. Moreover, in supplemental Figure S.1, we further investigated the stability of these motifs over time, and found that the usage difference between wt and tg is stable over the measured 25 minutes. These results confirm, demonstrated by a use case of APP /PS1 mice, that VAME can reveal behavioral differences that otherwise would remain undetected by human observation.

**Table 1:** Significant different motifs between tg and wt.

| Motif | Community | Mean Usage tg (%) | Mean Usage wt (%) | SE difference | T-ratio | p-Value |
| --- | --- | --- | --- | --- | --- | --- |
| 14 | Turning | 1.4 | 2.4 ↑ | 0.003 | 3.69 | 0.005 |
| 31 | Stationary | 2.6 | 3.4 ↑ | 0.007 | 2.827 | 0.008 |
| 34 | Walking | 1.4 ↑ | 0.9 | 0.004 | 3.839 | 0.0003 |
| 48 | Unsupported Rearing | 2.5 ↑ | 1.1 | 0.004 | 3.004 | 0.006 |
| 42 | Unsupported Rearing | 2.7 ↑ | 1.3 | 0.004 | 2.861 | 0.008 |

### 2.3 Motif transitions and behavioral dynamics

In the VAME framework, discrete representations of animal behavior are organized on different scales, varying from single motifs to communities. On the community level, the temporal structure of behavior can be identified by observing the probability of a community transitioning into another. The resulting transition matrices can be constructed both, on the community level or the motif level (see Methods 4.4). Figure 3 A shows the transition matrices for wt and tg animals ordered by the community structure. It can be seen that both groups share a similar structure of transitions, as expected given the similar open field behavior observed (Webster et al., 2013; Biallosterski et al., 2015; Giovannetti et al., 2018). To examine differences in transition probabilities between motifs we created a subtraction matrix 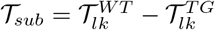 that illustrates, which transitions are more pronounced in wt animals (Red) or tg animals (Blue). indeed,we found significant differences in the usage of transitions within communities (Figure S.8, S.9). Overall, the most prominent differences in transition probabilities appeared in the *Stationary* community as well as the *Walking* community (Figure S.8).

**Figure 3:**
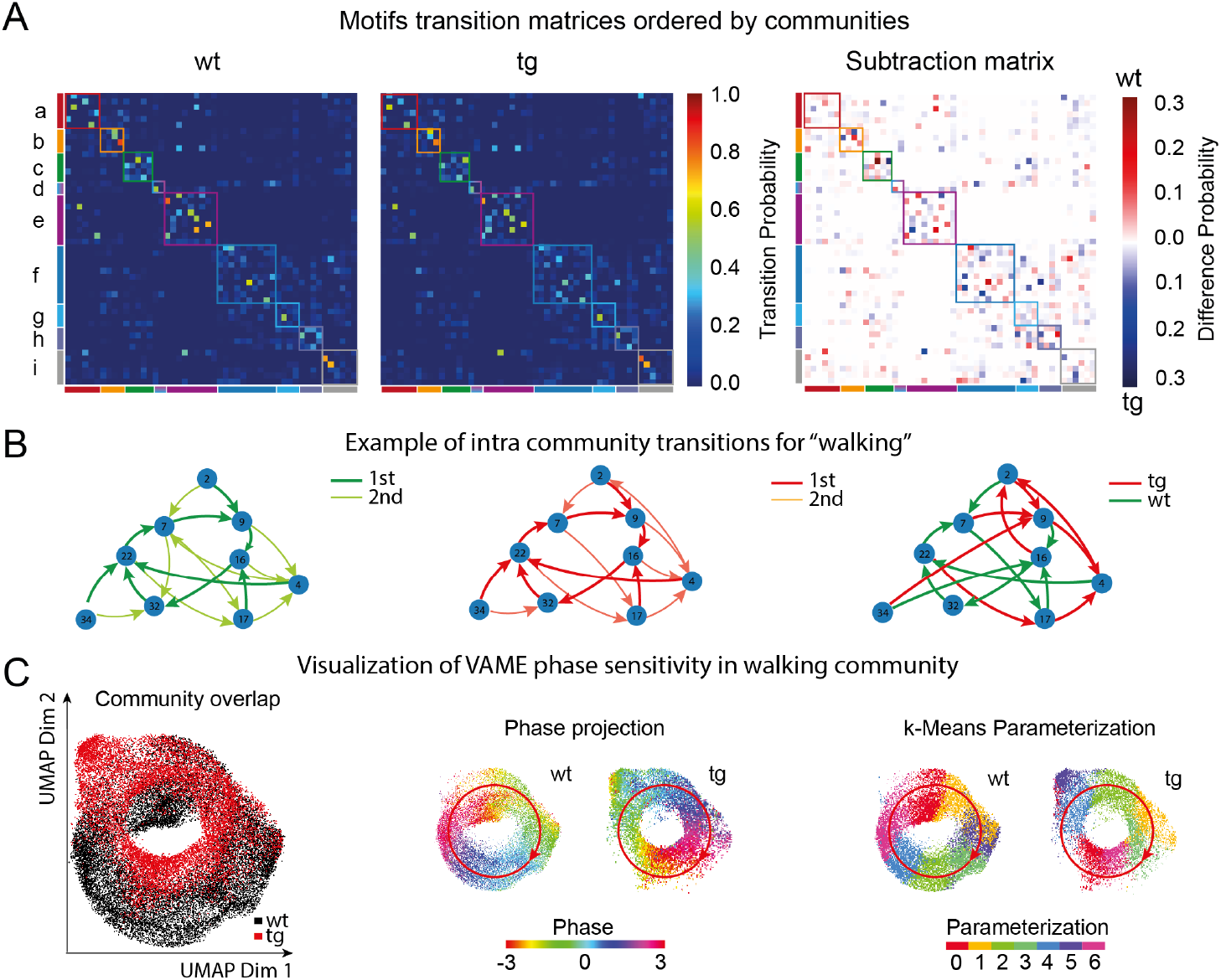
Identification of transition structure and locomotion patterns. (A) Left: Transition probability matrices ordered by communities for the wt and tg group. Right: Difference plot of both matrices. Squares along the diagonal indicate the grouping into communities. (B) Example of an intra-community transition graph for the walking community. We show the first and second highest transition for both groups (left, middle) and the highest transition difference (right). (C, left) Joint UMAP embedding of points belonging to the walking community in a wt (19.783 points, black) and a tg (13.264 points red) mouse reveals a circular structure. (C, middle) Projection of the mean phase angle of the horizontal hind paw movement onto the embedding displays the cyclic phase space of the walking movement in both animals. (C, right) Parametrization of both point clouds with k-Means shows blocks organized around the cyclic structure. Red arrow indicates the phase direction.

We investigated the *Walking* community in more detail to learn more about the transition differences. When following along the highest transitions on the Markov graph for this community (Figure 3 B, left, middle), a cyclic structure appears. Within this cyclic structure, different patterns of walking motifs are more strongly used by both experimental groups (Figure 3 B, right). To understand this structure, we embedded the encoded latent vectors of the *Walking* community onto a two-dimensional plane via UMAP. We visualized the UMAP in Figure 3 for two example animals from the wt and tg group. To determine if this structure is indeed cyclic, we decoded all points back to the original traces containing marker movements. Then, we computed the mean phase for the horizontal hind paw movement using Fourier transformation. We projected the phase angle back onto the embedding of latent vectors and observed that the phase angle follows the curve of the cyclic embedding (Figure 4 C, middle, red arrow shows phase direction). To parameterize the structure in both animals, we applied k-Means clustering. This yielded discrete clusters organized along the cyclic embedding (Figure 4, C, right). Such patterns are known to emerge from oscillatory dynamics modeled by RNNs (Chaudhuri, Gerçek, Pandey, Peyrache, & Fiete, 2019; Rubin et al., 2019). Cyclic representation of walking behavior was recently described in a *Drosophila* locomotion study (DeAngelis, Zavatone-Veth, & Clark, 2019) and here we confirm the existence of this representation also in rodent locomotion. We explore the possibility to detect locomotion subpatterns from this embedded dynamics in the supplemental section 5.9.

**Figure 4:**
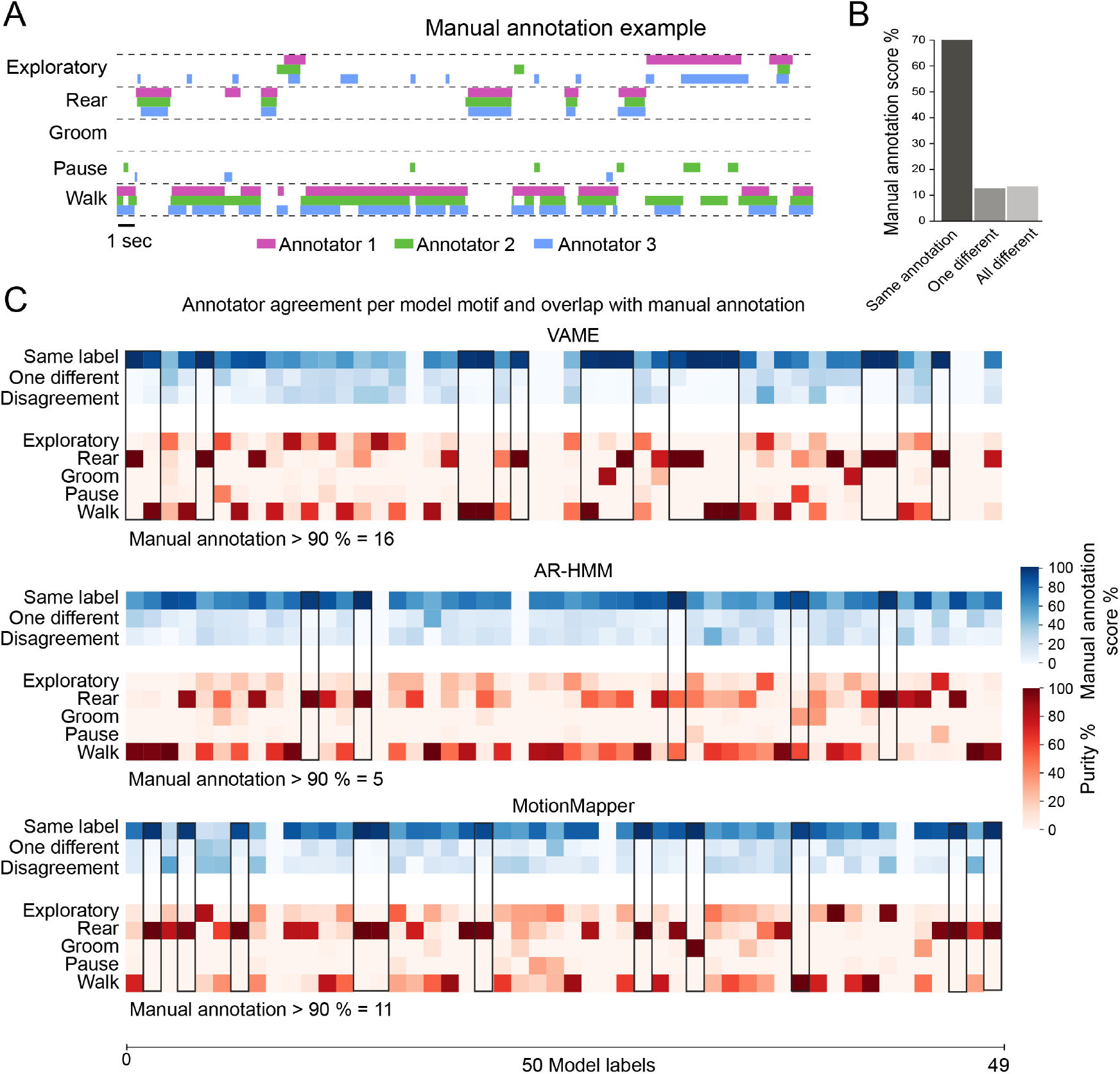
Annotated dataset and model comparison based on annotator agreement. (A) Overlap of manually assigned labels by three experts. (B) Disagreement in manual annotation. (C) Confusion matrices showing the annotator variability (blue) and the agreement between 50 model (VAME, AR-HMM, MotionMapper) motifs and 5 manually annotated labels (red). Empty columns exist when the specific motif did not appear in the annotated benchmark data.

### 2.4 Quantitative comparison of VAME with MotionMapper and AR-HMM

Several approaches for behavioral quantification exist, which all lead to valuable neuroethological data and provide important means for understanding the neural correlates of behavior in different model organisms (Datta et al., 2019). Since VAME makes use of the variational autoencoder framework, the approach differs substantially from others. Thus, we perfomed a qualitative (supplemental section 5.2 and 5.3) and quantitative comparison with two existing approaches to provide the user with useful information on which approach to preferably select in different experimental conditions and model systems.

In order to validate all three models (VAME, AR-HMM, MotionMapper) we created a manually labaled dataset that was annotated by three human experts with training in behavioral neuroscience. A video of a freely moving wt animal consisting of 20.000 frames (≈ 6 minutes length) was annotated with five behavioral labels (Walk, Pause, Groom, Rear, Exploratory behavior) (Figure 4 A, see Methods 4.5). When quantifying agreement between individual experts, we observed that 71.93% of the video frames were labeled equally by all three. The remaining 13.61% of frames were labeled unequally by two experts and 14.47% were labeled unequally by all three experts (Figure 4 B). This implicated that behavior showed a considerably high observer variability and is not trivially assignable to discrete labels (Anderson & Perona, 2014; Datta et al., 2019).

We trained all models on the full dataset and validated how they overlapped with the manual annotation (Figure 4 C). VAME had 16 motif overlapping with a high annotator agreement (> 90%) (Figure 4 C, black boxes). For MotionMapper, we identified 11 motifs which had a high annotator agreement and for the AR-HMM we identified 5 motifs with a high annotator agreement. This suggests that VAME is able to detect human identifiable labels in a more concise way than both other models.

To further investigate the overlap of each model with the benchmark dataset we quantified Purity, Normalized Mutual Information (NMI) and Homogeneity (see Methods 4.5). Applying all three measures we found that VAME had the highest score for each measure (Purity: 80.65%, NMI: 28.61%, Homogeneity: 54.89%) (Table 2) followed by the AR-HMM model. In order to rule out that this is an effect due to overclustering, we applied the same measurements also for smaller cluster sizes *K* =10 and *K* = 30, and could confirm that VAME achieved the highest scores on our benchmark (Table S.1). In supplemental Figure 4 C, we furthermore compared agreement overlap of the annotator with the model motifs. Here, we found that VAME shows the highest overlap followed by MotionMapper and AR-HMM. Moreover, in supplemental section 5.3 we investigate the spatiotemporal embedding of the VAME latent space by downprojecting it via Unifold Manifold Approximation (UMAP) and compare it to the MotionMapper t-SNE space.

**Table 2:** Quantitative model comparison based on an annotated benchmark dataset.

| k = 50 | Abs. Purity | Abs. NMI | Homogeneity % |
| --- | --- | --- | --- |
| MotionMapper | 70.67 | 17.35 | 30.82 |
| AR-HMM | 74.72 | 22.42 | 42.5 |
| VAME | **80.65** | **28.61** | **54.89** |

Our results indicate that, for the given approach of open field behavioral observation of mice, VAME showed a high degree of genralization to the dataset and identified behavioral motifs more robustly than others. However, while VAME is suitable for this approach and animal model, MotionMapper may be more suitable for the detection of motifs in fruit flies. More independent observations in different tasks and model systems would be required. Recently, data became available from an independent study on a head-fixed hand-reaching task in mice confirming our results and demonstrated the power of the autoregressive biRNN model in VAME compared to the AR-HMM model (Shi et al., 2021).

## 3 Discussion

In this manuscript, we introduce a new unsupervised deep learning method to discover spatiotemporal structure of behavioral signals obtained from pose estimation tools. Our method addresses a pressing need for behavior quantification in neuroscience, because current methods still insufficiently capture the complete spatiotemporal dynamics of behavior (Datta et al., 2019), and limit our understanding of its causal relationship with brain activity. It combines the VAE framework (Kingma & Welling, 2014) with autoregressive models (biRNNs). Thus, it creates a probabilistic mapping between the data distribution and a latent space. This approach enables the model to approximate the distribution and to encode factors that contain sufficient information to separate distinct behavioral motifs. A structural element of this model that differentiates it from other approaches is that it uses an additional biRNN decoder that learns to anticipate the structure of the subsequent trajectory and regularizes the latent space. This addition forces the encoder to learn a richer representation of the movement dynamics.

The field uses a rapidly developing repertoire of experimental approaches to investigate neural correlates of behavior (Zong et al., 2021; Xu et al., 2020; Steinmetz et al., 2021). Monitoring neural activity in freely behaving animals with imaging, electrophysiological tools and cell-type specific interrogation methods are state-of-the-art. In addition, new and traditional transgenic animal models allow for a deep investigation of molecular pathways in health and disease. All these approaches require deep, reliable and complete dissection of behavioral motifs. In this manuscript, we used a traditional transgenic model of Alzheimer’s disease, the APP/PS-1 model, to demonstrate discriminatory power of our approach in a mouse model system, which shows clear behavioral deficits in specific tasks, but no reported differences in open-field observation (Biallosterski et al., 2015; Giovannetti et al., 2018). Our approach, however, is not limited to any specific species or behavioral task as long as continuous video-monitoring can be provided (Luxem, Fuhrmann, Remy, & Bauer, 2019; Shi et al., 2021).

Our results demonstrate that VAME is capable of detecting differences between a tg and wt cohort of mice, while no differences were found by the human observer. In our use-case, we did not aim at investigating behavioral deficits in the domain of learning and memory with relation to Alzheimer’s disease. However, even in this small sample size, VAME identified a higher motif usage of the *Unsupported Rearing* community as well as a lower motif usage in the *Stationary* community that could be related to deficits of spatial orientation and environmental habituation (Lalonde et al., 2004; Janus et al., 2015).

We found that VAME is particularly suited to learn motif sequences from pose estimation signals. The main reason for this is that VAME is equipped with a particularly high sensitivity to the phase of the signals due to its biRNN encoder and decoder. In this manuscript, we showed this by plotting the phase angles onto a two-dimensional UMAP projection for the walking community. In this way, we could uncover a circularly organized point cloud, which exactly captured the natural cycle of limb movement (DeAngelis et al., 2019). Thus, VAME may be particularly useful in the detection of reoccurring locomotion patterns. While we so far have shown this only in one community, this advantage could be further exploited to identify differences in other movement types.

VAME is an unsupervised method that models the spatiotemporal information to quantify behavior, comparable to MoSeq and MotionMapper (Wiltschko et al., 2015; Berman et al., 2014). To incorporate spatiotemporal information, MoSeq applies an AR-HMM to infer hidden states from a series of transformed depth images. On the other hand, MotionMapper, which was initially implemented for fruit flies, relies on t-SNE (van der Maaten & Hinton, 2008) embeddings of wavelet transformations from high-speed camera images. In this method, regions with high densities are assumed to contain stereotypical behaviors. Since the spectral energy of a signal is the key input feature, low frequency movements, which are more prominent in mice than in flies, limits capturing the full behavioral repertoire. In contrast to MotionMapper, MoSeq was first applied in freely moving rodents. This allowed the detection sub-second behavioral structure but the AR-HMM resulted in a multitude of short and fast switching motifs, which can lead to uncertainty in animal action classification. These three methods have all been successfully applied to capture the behavioral dynamics in different animal models and experimental settings. To compare their performances, we trained all models on our data and investigated their motif sequence distribution on a benchmark dataset. Each model learned a consistent motif structure, nevertheless, VAME obtained the highest scores for all three metrics (Purity, NMI, Homogeneity). This result could be due to a better embedding of the spatiotemporal information and higher phase sensitivity of our model that is not as strongly present in the others. Recently, an independent group of scientists has published comparative data obtained from testing VAME to other approaches on a benchmark dataset for a hand-reaching task (Shi et al., 2021). Their findings fully support our own observation of higher VAME performance levels against other approaches (AR-HMM, Behavenet (Wiltschko et al., 2015; Batty et al., 2019)) in the following three criteria: Accuracy, NMI and Adjusted Rand Index. Indeed, the combination of the video representation model used in this study with VAME achieved the best results.

While VAME yielded higher performance scores when compared to other unsupervised approaches, supervised approaches may be better suited in experiments in which obtaining the full behavioral repertoire is not required. Supervised approaches like DeepEthogram, SimBa or MARS *DeepEthogram, SimBa or MARS* (Bohnslav et al., 2021; Nilsson et al., 2020; Segalin et al., 2021) all allow the labeling of episodes of interest. Lastly, *B-SOiD* (Hsu & Yttri, 2021), a recently developed unsupervised method that does not require a deep learning model, allows for a fast identification of similar frames and trains a classifier to identify these clusters rapidly in new data points. This approach, however, does not yield temporal information about the behavior and hence it will likely not capture the full range of behavioural dynamics. These aspects should be considered when choosing an optimal behavioral quantification method for a specific task and species. As VAME learns spatiotemporal information, it may be particularly useful to uncover behavioral dynamics. Moreover, VAME also has the potential to train a classifier on the latent vector information to quickly assign VAME motifs to new data points.

A promising future application of VAME could be the combination with three dimensional pose information (Günel et al., 2019; Sarkar et al., 2021; Dunn et al., 2021), which can be easily incorporated into the VAME model. This will likely lead to a better resolution of behavioral motifs, as most behaviors are expressed in three dimensions. When aiming at quantification of behavioral information with even higher dimensionality, the VAME model allows for a straightforward integration of parameters such as cellular calcium responses, neurotransmitter dynamics or other physiological features.

Taken together, we believe that VAME is an extremely useful new method for unsupervised behavior quantification, that can be easily applied by other scientists and strongly facilitate the investigation of causal relationships between brain activity and visible naturalistic behavior. VAME can identify motif structure from pose estimation signals with a high degree of generalization between animals and experimental setups. The framework is open-source and easily accessible without expert knowledge in programming and machine learning, and thereby open to a wide audience of neurobiologists and other scientists. We anticipate that it will stimulate the development of further machine learning models (Sun et al., 2021; Whiteway et al., 2021; Shi et al., 2021) and will trigger the development of robust metrics and benchmark datasets for the computational ethology community.

## 4 Methods

### 4.1 Animals

For all experiments we used 12 month old male transgenic and non-transgenic APPSwe/PS1dE9 (APP/PS1) mice (Jankowsky et al., 2001) on a C57BL/6J background (Jackson Laboratory). Mice were group housed under standard laboratory conditions with a 12-h light-dark cycle with food and water ad libitum. All experimental procedures were performed in accordance with institutional animal welfare guidelines and were approved by the state government of North Rhine-Westphalia, Germany.

### 4.2 Experimental setup, data acquisition and preprocessing

In the open field exploration experiment mice were placed in the center of an circular area (transparent plexiglas floor with diameter of 50 cm surrounded by a transparent plexiglas wall with height of 50 cm) and have been left to habituate for a duration of 25 minutes. Afterwards, sessions of 25 minutes were recorded where the mice were left to freely behave in the arena. To encourage a better coverage, three chocolate flakes were placed uniformly distributed in the central part of the arena prior to the experiment.

Mouse behavior was recorded at 60 frames per second by a CMOS camera (Basler acA2000-165umNIR) equipped with wide angle lens (CVO GM24514MCN, Stemmer Imaging) that was placed centrally 35 cm below the arena. Three infrared light sources (LIU780A, Thorlabs) were placed 70 cm away from the center, providing homogeneous illumination of the recording arena from below. All recordings were performed at dim room light conditions.

To extract behavioral pose, six virtual markers were placed on relevant bodyparts (nose, tailroot, paws) in 650 uniformly picked video frames from 14 videos. A residual neural network (ResNet-50) was trained to assign the virtual markers to every video frame (Mathis et al., 2018). The resulting training error was 2.14 pixels and the test error 2.51 pixels, respectively.

To obtain an egocentric time series of (*x, y*) marker coordinates we aligned the animal video frames from its allocentric arena coordinates to its egocentric coordinates. In order to get a tail to nose orientation from left to right we compute a rotation matrix and rotate the the resulting frame around the centre between nose and tail. This results into egocentrically aligned frames and marker coordinates **X** ∈ ℝ^*N×m*^, where *N* represents the recording length, i.e. 90000 frames and *m* the *x,y* marker coordinates. To fit our machine learning model we randomly sampled subsequences **x**_*i*_ from **X** that represent 500 ms of behavior i.e. 30 video frames. In the same manner, we created 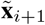 that stores the 15 subsequent time points of **x**_*i*_ to train the prediction decoder.

### 4.3 Variational Animal Motion Embedding

Given a set of *n* multivariate time series **X** = {**x**_1;_ **x**_2_,..., **x**_*n*_}, where each time series 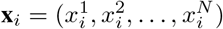 contains *N × m* ordered real values. The objective of our model is to learn a d-dimensional latent space **Z** by mapping the input **x**_*i*_ to a vector representation **z**_*i*_ ∈ ℝ^*d*^. Hence, **z**_*i*_ is learned via the non-linear mappings *f_enc_*: **x**_*i*_ → **z**_*i*_ and 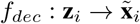, where *f_enc_, f_dec_* denotes the encoding and decoding process, respectively and is defined by,

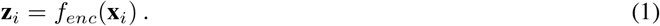

In order to learn the spatiotemporal latent representation our model encoder is parameterized by a two layer bi-directional RNN (biRNN) with parameters *ϕ*. Furthermore, our model uses two biRNN decoder with parameters *θ* and *η*.

The input data is temporally dependent and biRNNs are a natural choice in order to capture temporal dynamics. They extend the unidirectional RNN by introducing a second hidden layer which runs along the opposite temporal order. Therefore, the model is able to exploit information about the temporal structure from the past and future at the same time. Its hidden representation is determined by recursively processing each input and updating their internal state **h**_*t*_, at each timestep for the forward and backward path via,

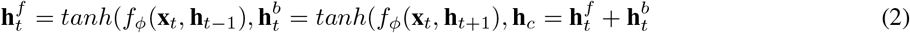

where 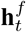 is the hidden information of the forward pass and 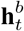 is the hidden information of the backward pass, *f* is a non-linear transition function, and *ϕ* is the parameter set of *f*. The transition function *f* is usually modelled as long short-term memory (LSTM) (Hochreiter & Schmidhuber, 1997) or gated recurrent unit (GRU) (Cho et al., 2014). Here, we use GRUs as transition function in the encoder and decoder.

The joint probability of a time series **x**_*i*_ is factorized by a RNN as product of conditionals,

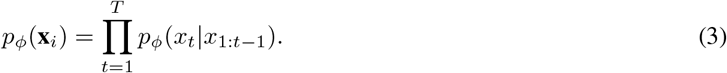

In order to learn a joint distribution over all variables, or more precise, the underlying generative process of the data, we apply the framework of variational autoencoders (VAE) introduced by (Kingma & Welling, 2014; Rezende et al., 2014). VAEs have been shown to effectively model complex multivariate distirbutions and can generalize much better across datasets.

#### 4.3.1 Variational Autoencoder

In brief, by introducing a set of latent random variables **Z** the VAE model is able to learn variations in the observed data and can generate **X** through conditioning on **Z**. Hence, the joint probability distribution is defined as,

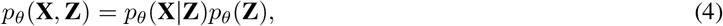

parameterized by *θ*.

Determining the data distribution *p*(**X**) by marginalization is intractable due to the non-linear mappings between **X** and **Z** and the integration of **Z**. In order to overcome the problem of intractable posteriors the VAE framework introduces an approximation of the posterior *q_ϕ_*(**Z**|**X**) and optimizes a lower-bound on the marginal likelihood,

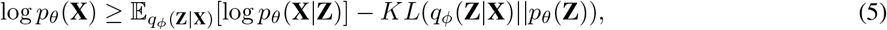

where *KL*(*Q*||*P*) denotes the Kullback-Leibler divergence between two probability distributions *Q* and *P*. The prior *p_θ_* (**Z**) and the approximate posterior *q_ϕ_*(**Z**|**X**) are typically chosen to be in a simple parametric form, such as a Gaussian distribution with diagonal covariance. The generative model *p_θ_*(**X**|**Z**) and the inference model *q_ϕ_*(**Z**|**X**) are trained jointly by optimzing Eq. 5 w.r.t their parameters. Using the *reparameterization trick* (Eq. 6), introduced by (Kingma & Welling, 2014) the whole model can be trained through standard backpropagation techniques for stochastic gradient descent.

#### 4.3.2 Variational lower bound of VAME

In our case, the inference model (or encoder) *q_ϕ_*(**z**_*i*_|**x**_*i*_) is parameterized by a biRNN. By concatenating the last hidden states for the forward and backward path we obtain a global hidden state **h**_*i*_, which is a fixed-length vector representation of the entire sequence **x**_*i*_. To get the probabilistic latent representation **z**_*i*_ we define a prior distribution over the latent variables *p_θ_*(**z**_*i*_) as an isotropic multivariate Normal distribution 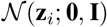. Its parameter *μ_z_* and Σ_*z*_ of the approximate posterior distribution *q_ϕ_*(**z**_*i*_|**x**_*i*_) are generated from the final encoder hidden state by using two fully connected linear layers. The latent representation **z**_*i*_ is then sampled from the approximate posterior and computed via the reparameterization trick,

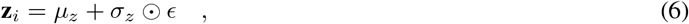

where *ϵ* is an auxiliary noise variable and ⊙ denotes the Hadamard product.

The generative model *p_θ_* (**x**_*i*_ |**z**_*i*_) (or decoder) receives **z**_*i*_ as input at each timestep *t* and aims to reconstruct **x**_*i*_. We use the mean squared error (MSE) as reconstruction loss, defined by,

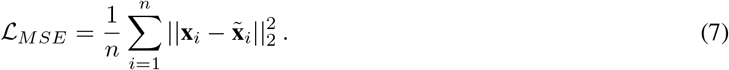

The log-likelihood of **x**_*i*_ can be expressed as in Eq. 5. Since the KL divergence is non-negative the log-likelihhood can be written as

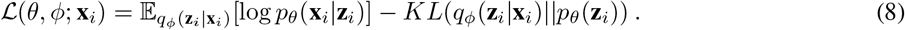

Here, 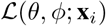 is a lower bound on the log-likelihood and therefore called the *evidence lower bound* (ELBO) as formulated by (Kingma & Welling, 2014).

We extend the ELBO in our model by an additional prediction decoder biRNN 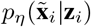 to predict the evolution 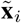 of **x**_*i*_, parameterized by *η*. The motivation for this additional model is based on (Srivastava et al., 2015), where the authors propose a composite RNN model which aims to jointly learn important features for reconstruction and predicting subsequent video frames. Here, 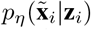 serves as a regularization for learning **z**_*i*_ so that the latent representation not only memorizes an input time series but also estimates its future dynamics. Thus, we extend Eq. 8 by an additonal term and parameter,

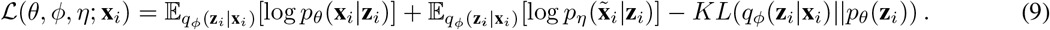

Finally, the training objective is to minimize

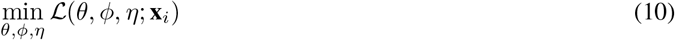

and the overall loss function can be written as

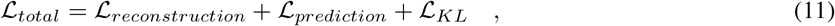

where 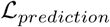 is the MSE loss of the prediction decoder.

The full model was trained on the combined dataset (1.3*e*6 time points) using the Adam optimizer (Kingma & Ba, 2015) with a fixed learning rate of 0.0005 on a single Nvidia 1080ti GPU. All computing was done with PyTorch (Paszke et al., 2017). The ergodic mean of the reconstruction error 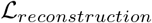 for all virtual marker time series was found to be 1.82 pixels.

### 4.4 Motif identification

To determine the set of behavioral motifs *B* = {*b*_1_,..., *b_K_*} we obtained the latent vector **Z** from a given dataset using VAME as described in Methods 4.3. Here, **Z** ∈ ℝ^*d×N–T*^, with the embedding dimension d, the number of frames *N* and the temporal window *T*, represents the feature space from which we want to identify motifs. By treating the underlying dynamical system as a discrete-state continuous-time Markov chain, we apply a Hidden-Markov-Model (HMM) with a Gaussian distributed emission probability to this space to detect states (motifs). We used the **hmmlearn** python package in our framework to implement the HMM. The default settings for the Gaussian emission model from the packages were used. Moreover, we compared the HMM to a simpler and less time consuming k-means clustering in supplemental section 5.4. To identify the number of motif present in our dataset, we used a similar approach as in (Wiltschko et al., 2015). We let a HMM infer 100 motifs and identified the threshold, where motif usage dropped below 1%, see supplemental Figure S.3. Motif usage was determined as the percentage of video frames that are assigned to the occurrence of a specific motif.

To model the transitions between behavioral motifs, we interpreted the motif sequence as a discrete-time Markov chain where the transition probability into a future motif is only dependent on the present motif. This results in a *K × K* transition probability matrix 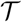, with the elements

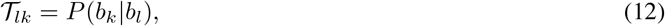

being the transition probabilities from one motif *b_l_* ∈ *B* to another motif *b_k_* ∈ *B*, that are empirically estimated from clustering of **Z**.

In order to obtain a hierarchical representation of behavioral motifs we can represent the Markov chain (12) as a directed graph 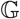 consisting of nodes *v*_1_...*v_K_* connected by edges with an assigned transition probability 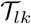 We can transform 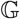 into a binary tree 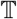 by iteratively merging two nodes (*v_i_, v_j_*) until only the root node *v_R_* is left. Every leaf of this tree represents a behavioral motif. To select *i* and *j* in each reduction step, we compute the cost function

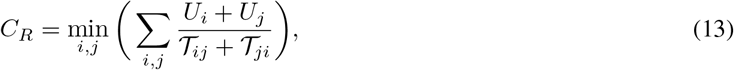

where *U_i_* is the prooaomty or occurrence tor the *i*th motif. Note that after each reduction step the matrix 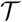 is recomputed in order to account for the merging of nodes. Lastly, we obtain *communities* of behavioral motifs by cutting 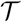 at given depth of the tree, analogous to the hierarchical clustering approach used for dendrograms.

### 4.5 Manually assigned labels and scoring

In order to obtain manually assigned labels of behavioral motifs we asked three experts to annotate one recording of freely moving behavior with a duration of 6 minutes. All three experts had a strong experience with in-vivo experiments as well as ethogram-based behavior quantification. The experts could scroll trough the video in slow-motion forward and backward in time and annotated the behavior into several atomic motifs as well as a composition of those. As an example, the experts were allowed to annotate a behavioral sequence as *walk* or *exploration*, but also *walk and exploration*. We then summarized the annotation into atomic motifs into 5 coarse behavioral labels, as shown in Table 3.

**Table 3:**
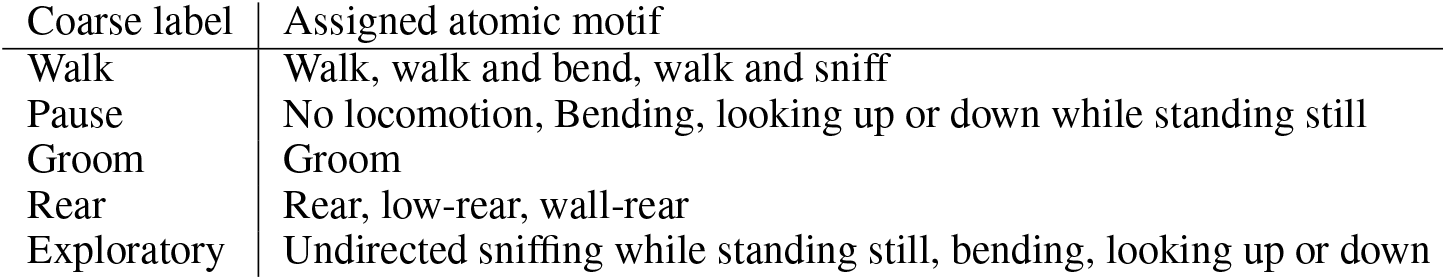
Assignment of atomic motifs into coarse behavior labels.

The coarse labels were created with respect to the behavior descriptions taken from the Mouse Ethogram database (www.mousebehavior.org), which provides a consensus of several previously published ethograms. The assignment of coarse labels to the Mouse Ethogram database taxonomy is shown in Table 4.

**Table 4:** Mouse Ethology database taxonomy corresponding for each manually assigned coarse label.

| Coarse label | Mouse Ethogram database |
| --- | --- |
| Walk | Active behavior - General activity - Exploratory behavior - Search - General locomotion |
| Pause | Inactive behavior- Still and alert |
| Groom | Active behavior - Maintenance behaviors - Grooming |
| Rear | Active behavior - General activity - Exploratory behavior - Search - Rearing |
| Exploratory | Active behavior - General activity - Exploratory behavior - Investigate - Undirected sniffing |

For scoring of human assigned labels to the behavioral to VAME motifs we used the clustering evaluation measures Purity, NMI and Homogeneity. Purity is defined as

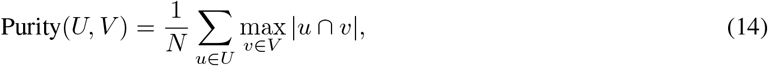

where *U* it the set of manually assigned labels *u, V* is the set of labels generated by VAME *v* and *N* is the number of frames in the behavioral video. The Normalized Mutual Information score is written as

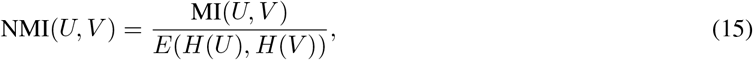

where MI(*U, V*) is the mutual information between set *U* and *V* defined as

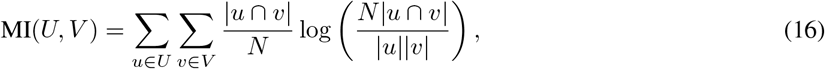

and *H*(*U*) is the entropy of set *U* defined as

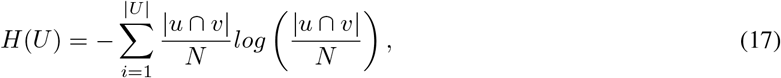

where the || operator denotes the amount of frames that have the corresponding labels assigned. Homogeneity is defined as

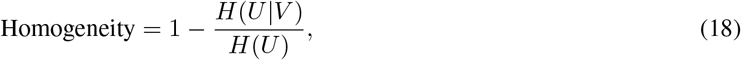

where the conditional entropy of manually assigned labels given the cluster assignments from VAME is given by

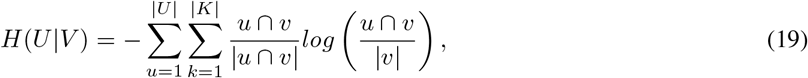

Note that the Purity score (14) is larger when the set *V* is larger than *U* and the NMI score (15) is generally larger when both sets *U* and *V* are of similar size, i.e. the number of possible labels is roughly the same in the human assigned set as well as the set generated using VAME.

### 4.6 Human phenotype classification task

For the classification of phenotypes using human experts we have created an online form, where experts could watch all eight videos and make their choice about which phenotype is shown in each video. There was no time limit and the average time to complete the questionnaire was 30 minutes. The participants have not been told how many animals of each group are in the set. For every video, the following five decision could be made: APP/PS1 (Very sure), APP/PS1 (Likely), Unsure, Wildtype (Likely), Wildtype (Very Sure). We have counted a right answers *(Very sure* and *Likely)* as a correct classification (1 point), and wrong answers as well as the choice for the *Unsure* option as wrong classification (0 points). Eleven experts were participating in this classification task. All of them had previous experience with behavioral video recordings in an open field and/or treadmill setting. In addition, six of the participants had previous experience with the APP/PS1 phenotype.

### 4.7 Code availability

The VAME toolbox is available at https://github.com/LINCellularNeuroscience/VAME.

## 5 Acknowledgments

We thank J. Macke, E. Restrepo, J. Gall and S. Stober for comments on the manuscript. This work was supported by the European Research Council (CoG;SUBDECODE) and DFG-SFB 1089.

## Supplemental Materials

### 5.1 Motif description and time binned evolution

**Figure S.1:**
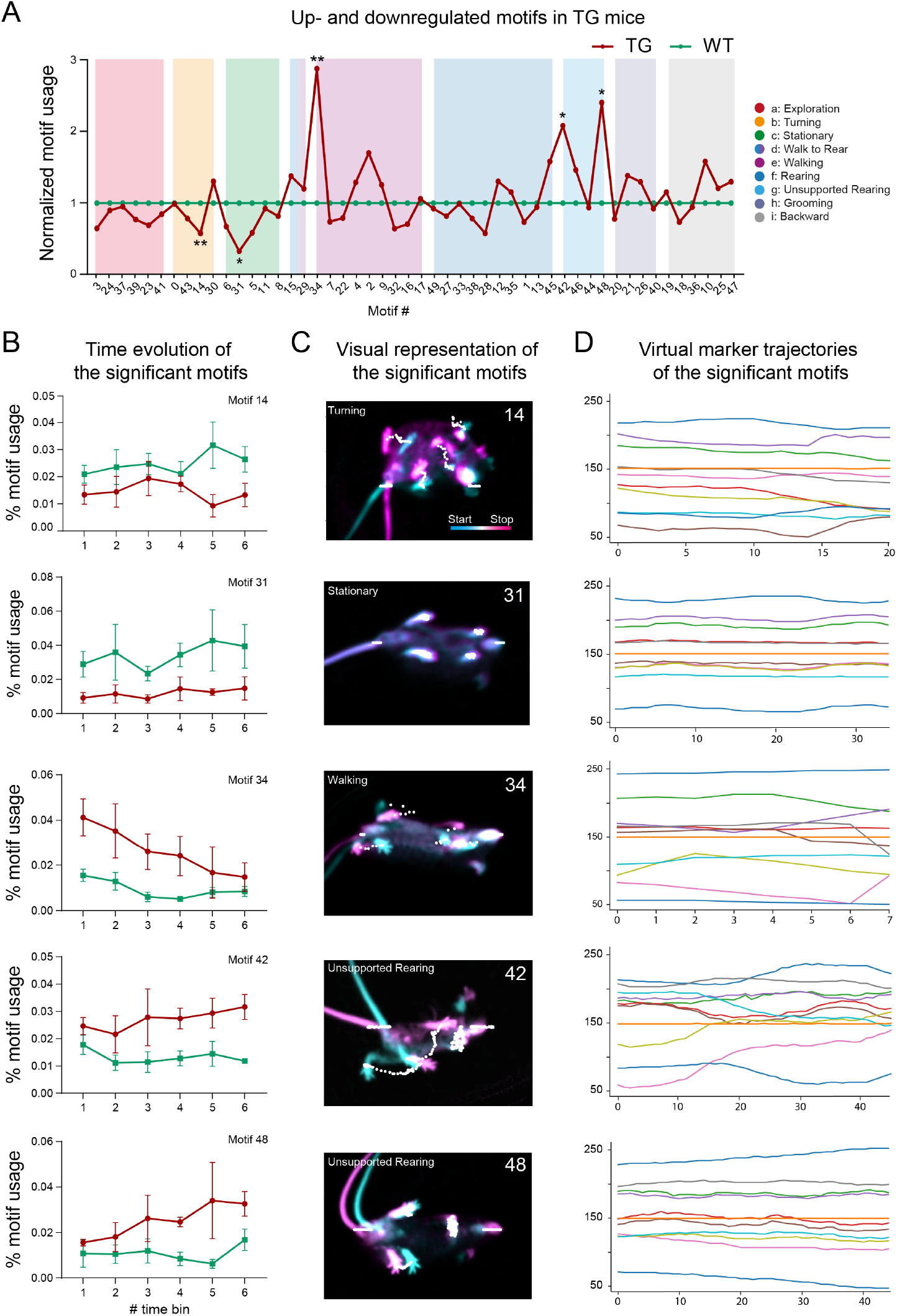
Phenotype specific and time dependent modulation of behavioral motifs. (A) Depiction of up- and downregulation of motifs and communities in TG animals (red line) compared to WT animals (green line) and ordered by communities. (B) Binned time evolution of significant motifs between both phenotypes. (C) Visual representation of significant motifs. The start frame is colored in cyan and the end frame is colored in magenta. White dots represent the DLC virtual marker points. (D) Corresponding exemplary (x,y)-virtual marker trajectory of the visual representation shown in C.

Figure S.1 A shows an alternative way to represent behavioral differences between two groups or phenotypes. Here, the usage of a motif for the tg group is calculated as a ratio and normalized against the wt group. This representation depicts the up- or downregulation in motifs or communities with respect to the tg group. The *Exploration, Turning* and *Stationary* communities are downregulated, while the *Walk to Rear* and *Unsupported Rearing* communities are upregulated. In the *Walking* community, the biggest peak of upregulation happens in the significant changed motif 34. Other communities have certain motifs which are differently used but with no significant group difference.

In order to investigate how stable the significant differences are during the experiment, we binned the experiment into six equal blocks (Figure S.1 B). We find that the motif usage is constantly up- or downregulated for all significantly changed motifs. In Figure S.1 C we visualized the these motifs by taking the start (cyan color) and end (magenta color) frame for a random episode of motif occurrence. White dots are representing the DeepLabCut virtual marker positions over time. Panel D of Figure S.1 shows the corresponding DeepLabCut trajectory of the visualized motif.

To describe the motifs we investigated the single motif videos (Examples are given in supplemental video 1-5). Motif 14 *(Turning* community) is characterized by a rotational motion with the front paws further away from each other. Motif 31 *(Stationary* community) shows no paw movement but some nose motion. Motif 34 ( *Walking* community) shows a push into a slow walk motion and as shown on the hierarchical tree Figure 2, C, it is grouped further away from the other walk motifs, which suggests that it does not belong to the classical walk cycle. Motif 42 and Motif 48 *(Unsupported Rearing* community) show a stationary standing animal inside the arena with some nose motion. Stationary here refers to almost no movement in the back paws.

### 5.2 Model comparison

**Figure S.2:**
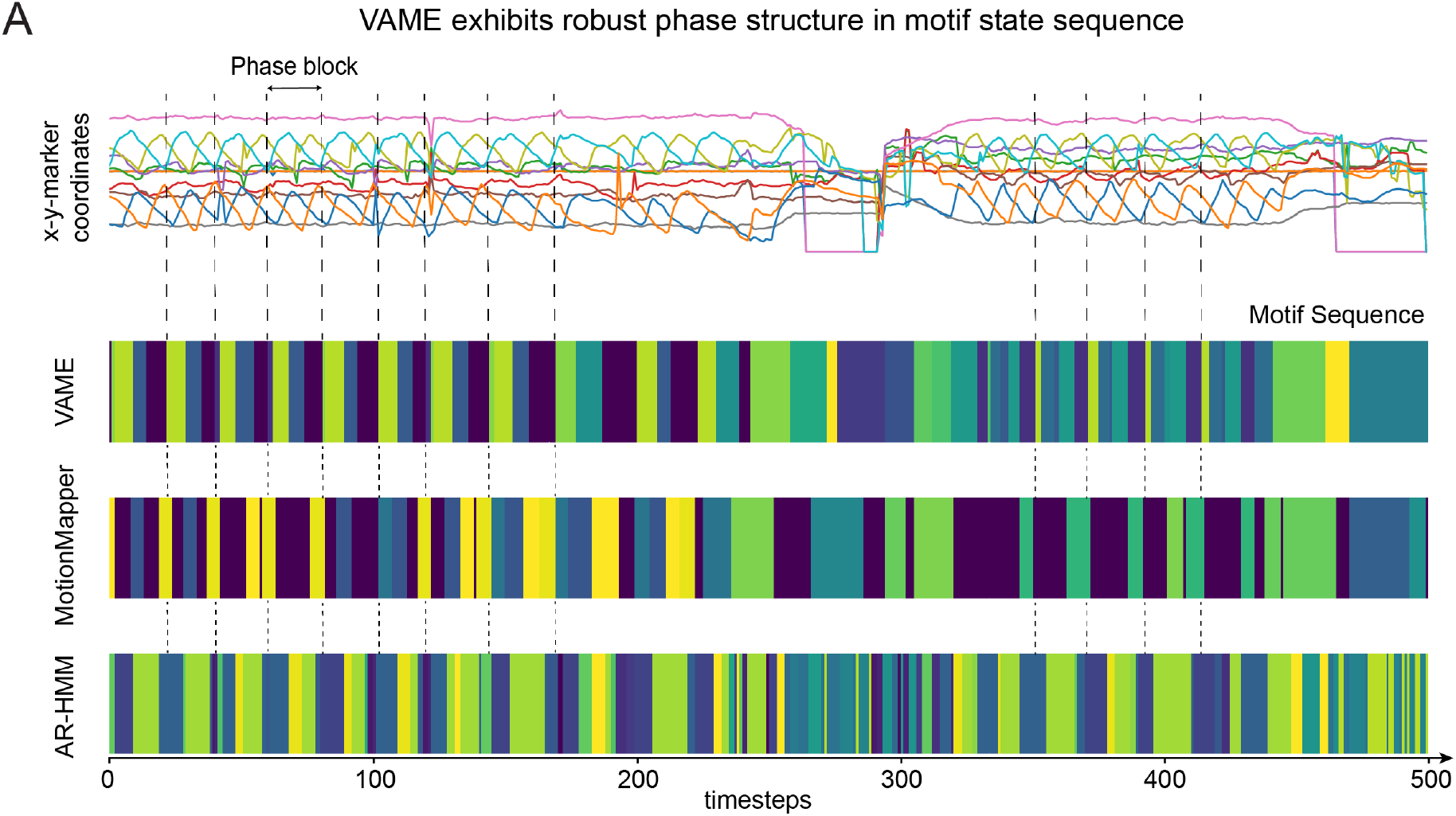
Qualitative comparison with MotionMapper and AR-HMM. (A) Exemplary trace of the input time series together with the motif segmentations obtained from the VAME, MotionMapper and AR-HMM methods.

Figure S.2 A shows a qualitative comparison between VAME and current methods (MotionMapper and AR-HMM, (Berman et al., 2014; Wiltschko et al., 2015)). We trained all models on our data and carried out segmentation into approximately the same amount of behavioral motifs (VAME, AR-HMM n=50, MotionMapper n=51). The exemplary trace is aligned to the motif sequences (Figure 4, top) and shows two walking and rearing episodes. As reference, we selected the x-coordinate of the left hind paw (orange marker) to define a phase block, indicated by the dashed lines between time step 0 and 200 (Figure 4, top). Visual inspection between the methods revealed that VAME motifs match the phase of the signal more precisely. The motif sequence obtained from AR-HMM exhibits more rapid state switches while MotionMapper has more longer lasting motifs.

Table S.1 reports the results on Purity, NMI and Homogeneity for smaller cluster sizes of *k* = {10,30} for all three models (VAME, MotionMapper, AR-HMM) on our benchmark dataset.

**Table S.1:** Model comparison based on an annotated dataset with *k* =10 and *k* = 30 motifs.

| Model | Purity | NMI | Homogeneity % |
| --- | --- | --- | --- |
| MotionMapper ($k = 10$) | 62.14 | 10.91 | 15.14 |
| AR-HMM ($k = 10$) | 68.12 | 18.69 | 26.15 |
| VAME ($k = 10$) | **74.20** | **29.65** | **42.00** |
| MotionMapper ($k = 30$) | 66.37 | 14.13 | 22.98 |
| AR-HMM ($k = 30$) | 73.99 | 21.77 | 37.97 |
| VAME ($k = 30$) | **79.19** | **26.10** | **51.16** |

### 5.3 Downprojection and trajectories

**Figure S.3:**
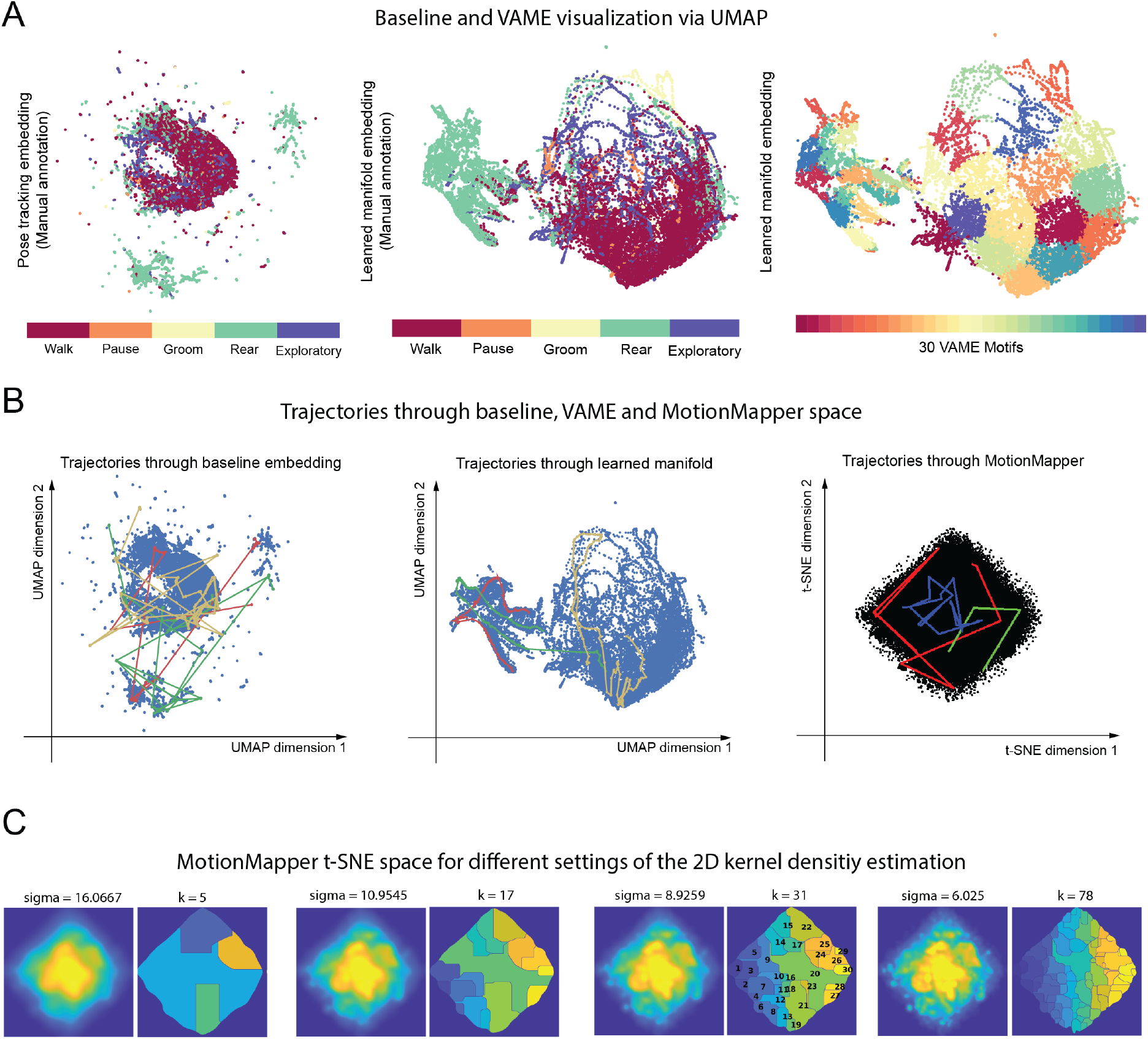
Model embedding and trajectories. (A) Unifold Manifold Approximation (UMAP) embeddings of the marker position input series (left), the latent representation encoded from the RNN color-coded for 5 manually labeled behavioral classes (middle) and color-coded according to assignment into 30 VAME motifs (right). (B) Three exemplary paths of consecutive video frames through the UMAP embedding space of the spatial input series (left) and through the embedding space visualizing the spatiotemporal representation (Middle). (Right) t-SNE embedding obtained by MotionMapper with three examplary paths. (C) Embeddings and segmentations obtained from MotionMapper for different settings of the 2D kernel density estimation parameter (sigma).

In Figure S.3 A (left) we show the Unifold Manifold Approximation (UMAP) projection of the original egocentrically aligned virtual marker signal (baseline) for the manual annotated dataset and (middle, right) the UMAP projection of the latent vectors obtained by VAME for the same dataset (middle: human label, right: VAME motifs). It can be observed that the VAME embedding is less scattered and more densely represented.

To test the spatiotemporal connectivity of the latent representation for VAME, we sampled three random behavioral sequences with a length of 1.5 seconds and plotted their trajectories on top of both visualizations. Moreover, we also plotted this trajectory on top of the t-SNE embedding of MotionMapper as comparison. We observed that the course of trajectories within the embedding of VAME followed a smooth path through the projected manifold while trajectories through the embedding of the baseline signal appears scattered. For MotionMapper, we also observed more scatter (Figure S.3 B).

Lastly, we investigated the embedding of MotionMapper more closely to understand the model behavior on our dataset (Figure S.3, C). More specifically, we show the effect of modifying the standard deviation of the smoothing Gaussian of the 2D kernel density estimation, a free parameter of MotionMapper, on the obtained t-SNE embedding as well as on obtained the watershed segmentations. As depicted here, the choice of the parameter has a strong impact on the clusterability; a high setting of the standard deviation implies a lower number of clusters that are detectable by the watershed segmentation, and vice-versa. Here, we observe that the overrepresented density in the middle of the embedding exists for segmentations to 5, 17 and 31 clusters, and is only resolved for a large setting of the cluster size (k=78). This suggests that the largest mass of the behavioral space is mostly contained in a single motif located in the center of the space. As discussed previously, this finding is likely observable for pose estimation data obtained from mice, as the sinusoidal signals have a higher similarity in the frequency space, when compared to behavioral signals measured for, for example, fruit flies.

### 5.4 VAME model selection

The VAME model consists of one biRNN encoder and two biRNN decoder. A HMM is used to identify hidden states (motifs) within our embedding space, as described in section 2.1 (also see Methods 4.3). The model was chosen after we tested four different variations of the architecture and compared the HMM against a k-Means algorith. In table S.2, we show the different choices and validated their outcome based on our benchmark dataset.

Our architectural choices were either a standard variational autoencoder consisting of a biRNN encoder and a biRNN decoder or with an additional biRNN prediction decoder. Furthermore, we applied to both variants spectral regularization of the latent space (Ma et al., 2019) to see if this could lead to improved clusterability. We applied three metrics (Purity, NMI, Homogeneity, see section 2.4) to identify the best model. In both cases (k-Means or HMM), the variational autoencoder model without spectral regularization and an additional decoder had the highest scores. The model with HMM led to the best scores and hence, we chose this as the primary model in our manuscript.

**Table S.2:** VAME model selection with two different segmentation algorithm (k-Means and HMM) for *k* = 50. Reported is the mean of five repeated training and inference runs.

| Model (segmentation) | Purity | NMI | Homogeneity % |
| --- | --- | --- | --- |
| VAME single decoder (k-Means) | 74.66 | 22.26 | 43.02 |
| VAME single decoder + spectral regularization (k-Means) | 76.17 | 22.13 | 43.20 |
| VAME two decoder (k-Means) | **76.23** | **23.58** | **45.55** |
| VAME two decoder + spectral regularization (k-Means) | 75.56 | 23.12 | 44.69 |
| VAME single decoder (HMM) | 78.44 | 26.70 | 49.82 |
| VAME single decoder + spectral regularization (HMM) | 79.81 | 26.98 | 51.98 |
| VAME two Decoder (HMM) | **80.66** | **28.61** | **54.85** |
| VAME two Decoder + spectral regularization (HMM) | 79.15 | 27.64 | 52.84 |

### 5.5 Motif clustering number

Deciding the number of cluster (e.g. motifs) *k* present in a data set is hard task when no a-priori knowledge is available. To determine this number for our dataset, we used a similar technique as presented in (Wiltschko et al., 2015). We took our trained VAME model and let the HMM infer 100 motifs and treated motifs that had less than 1% usage as noise. This mark was 50 motifs, which we used throughout the paper. Figure S.4 visualizes the motif usage for all eight animals (blue lines), the mean usage (red line) and the 1% threshold (dashed line). The first 10 motifs have a high usage, which then drops slowly to the 1% threshold line (48 motifs) and continues to decreae to almost 0% usage (100 motifs).

**Figure S.4:**
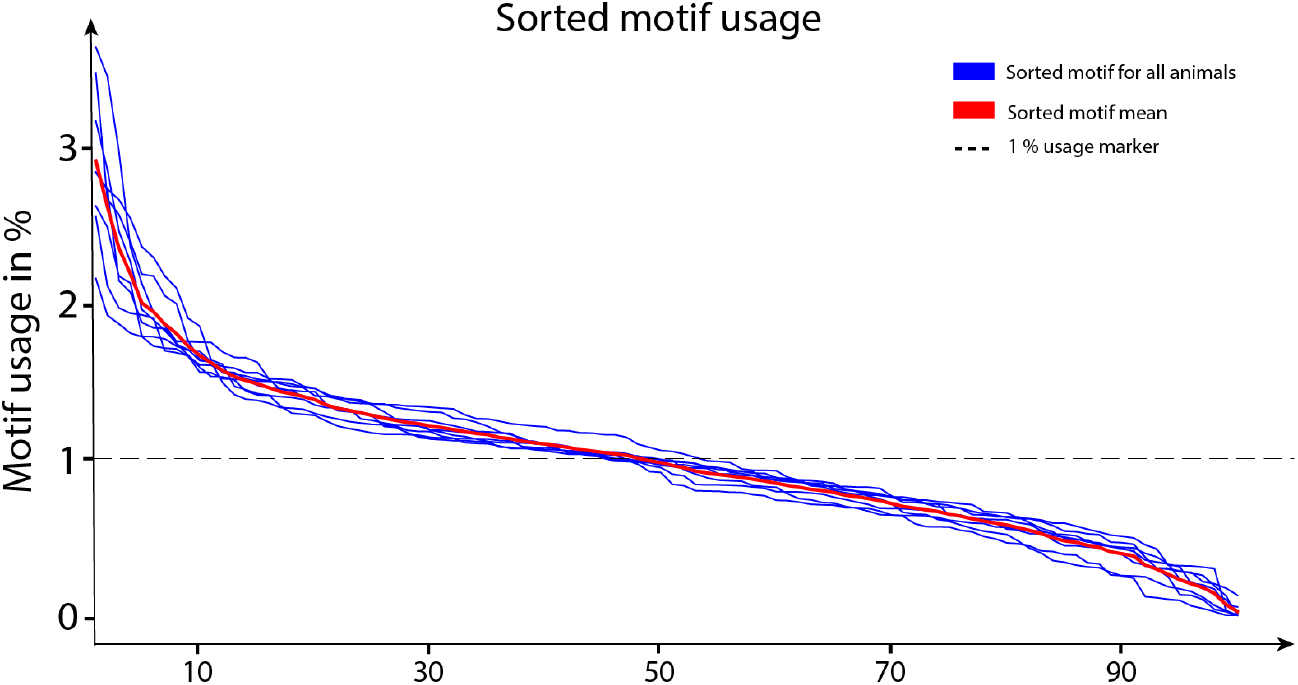
Sorted motif usage for all animals.

### 5.6 Parameters and biases

**Table S.3:** Table of choices for our approach and settings for our model.

| Choice | Our setting |
| --- | --- |
| What is the pixel resolution of the animal body in the camera image | In our case, the animal body was covered with approximately $100 \times 300$ pixels. |
| What is the temporal resolution of the camera? | The employed camera operates at 60 Hz. |
| How many virtual markers are used and how are they placed on the animal body parts? | We placed 6 virtual markers on paws, the nose and the tail root (see Figure 1). |
| Which software is used for keypoint detection? | We used DeepLabCut 1.1 (Mathis et al., 2018). |
| How many frames are manually labeled? | We labeled the keypoints in 600 uniformly chosen frames. |
| How long is the keypoint detection CNN trained? | We trained the keypoint detection for 350.000 iterations, the resulting training errors was 2.14 pixels and the test error was 2.51 pixels. |
| How is the obtained keypoint time series aligned to egocentric coordinates? | We use the procedure defined in the Methods section. |
| How is missing data handled? | Missing data points for periods where pose estimation could not reliably detect the position of the corresponding virtual marker shorter then 200 ms were linearly interpolated while the values for longer periods were set to an arbitrary negative value. |
| Which size of the time window was used for the reconstruction and prediction decoder? | We used 30 frames (500 ms) for the reconstruction and 10 frames (166 ms) for the prediction decoder. |
| Which other hyperparameters have been used to train VAME? | All other hyperparameter settings can be found in the default configuration file available at the project website. |
| How many datapoints are used to train VAME? | We trained our model with 1.3 million data points. |
| How is the clustering carried out? | We used a Hidden Markov Model for state detection as described in the Methods section. |

Although the proposed quantification method is unsupervised, the choice of parameters as well as processing steps integrate biases into the quantification result. In Table S.3 we summarize the biases of our approach and state the parameter setting that have been used for our data.

### 5.7 Community visualization and description

**Figure S.5:**
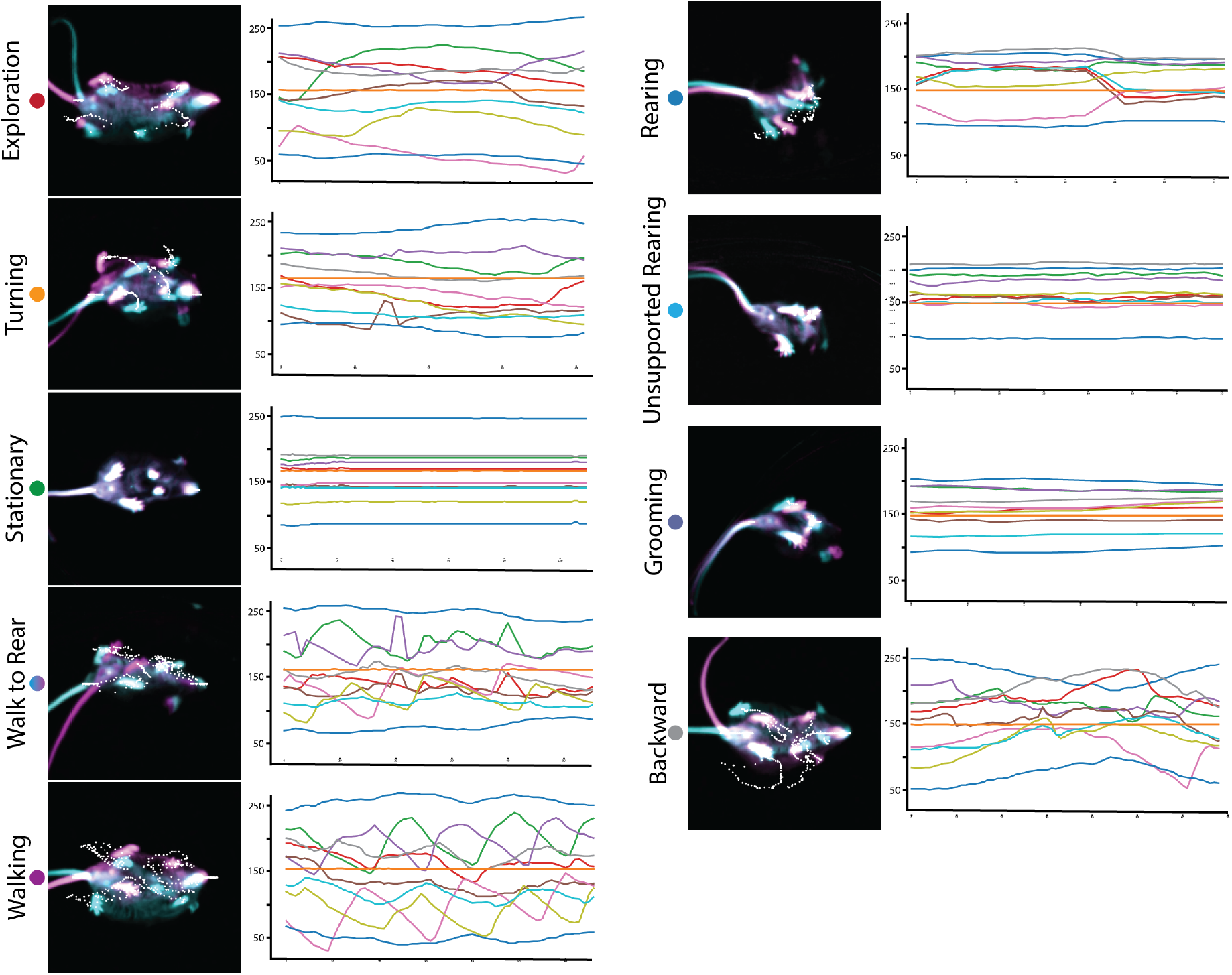
Visualization of Communities with their respective DLC trace.

In Figure S.5 we visualized all nine communities by taking the start (cyan color) and end (magenta color) frame for a random community episode. White dots are representing DeepLabCut marker. Next to the visual representation, the DeepLabCut trace for this episode is shown. Community *a* contains motifs with exploration characteristics such as slow walking and a lot of nose movement which could be interpreted as sniffing. Community *b* shows mainly events in which motifs express rotational behavior. In *c,* the motifs display almost no movement of any body part. Community *d* consists of two motifs which depict transitional behavior from walk to rear or vice versa. In community *e*, we found that all motifs express a specific part of the walking behavior. Community *f* contains motifs which are mainly showing rears along the wall of the arena while *g* contains motifs depicting rears within the arena. Community *h* belongs to the same branch as *g* but portrays mainly motifs with grooming activity. Lastly, community *i* shows motifs in which the animal performs a backward motion e.g after rearing.

### 5.8 Generative model aspects

**Figure S.6:**
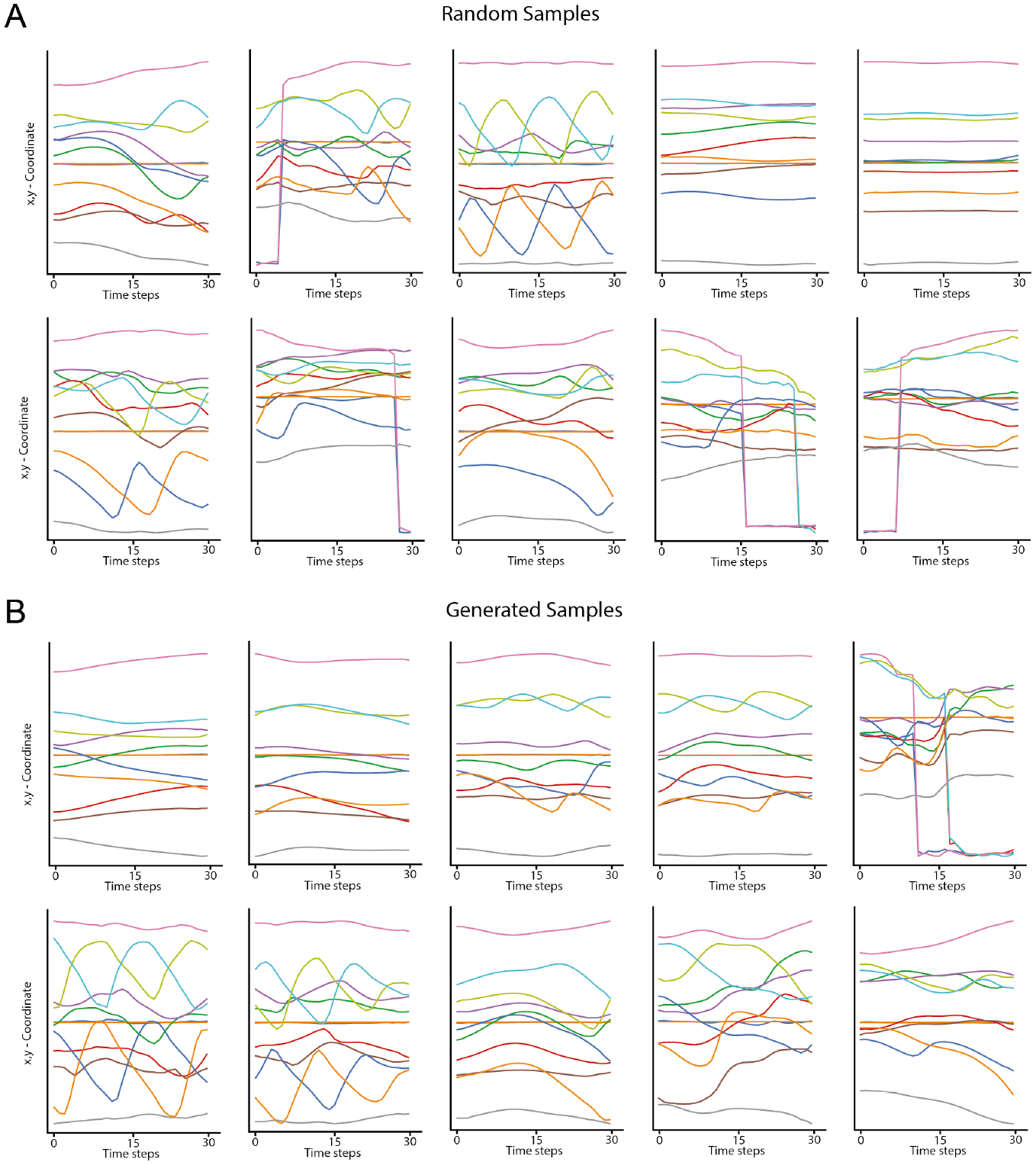
Random sample reconstruction using generative modeling. (A) Reconstruction of randomly drawn samples from the input data. (B) Generated samples obtained by fitting a Gaussian mixture model on the latent space, and decoding points sampled from the fitted density via the reconstruction decoder.

The VAME framework is based on the concept of variational autoencoder, which belongs to the class of generative models. The power of a generative models lies in its ability to learn the data distribution and create new, unseen samples from this distribution. This capability can be used to show that the model is able to represent the data distribution. In Figure S.6 A, we first show VAME’s capability to reconstruct random samples form the input data. To demonstrate that we can generate accurate samples from VAME’s latent space, we carried out ex-post density estimation using a 10-dimensional GMM, as proposed in (Ghosh, Sajjadi, Vergari, Black, & Scholkopf, 2020). From the fitted density, we are able to sample datapoints that can be transformed to input-traces using the decoder of the trained model. With this functionality, users of VAME can verify if the model has succesfully learned the data distribution to generate realistic synthetic samples. Note that this approach can be also used to validate single clusters, if the density is fitted on regions belonging to individual motifs only.

### 5.9 Sequence detection in locomotion

**Figure S.7:**
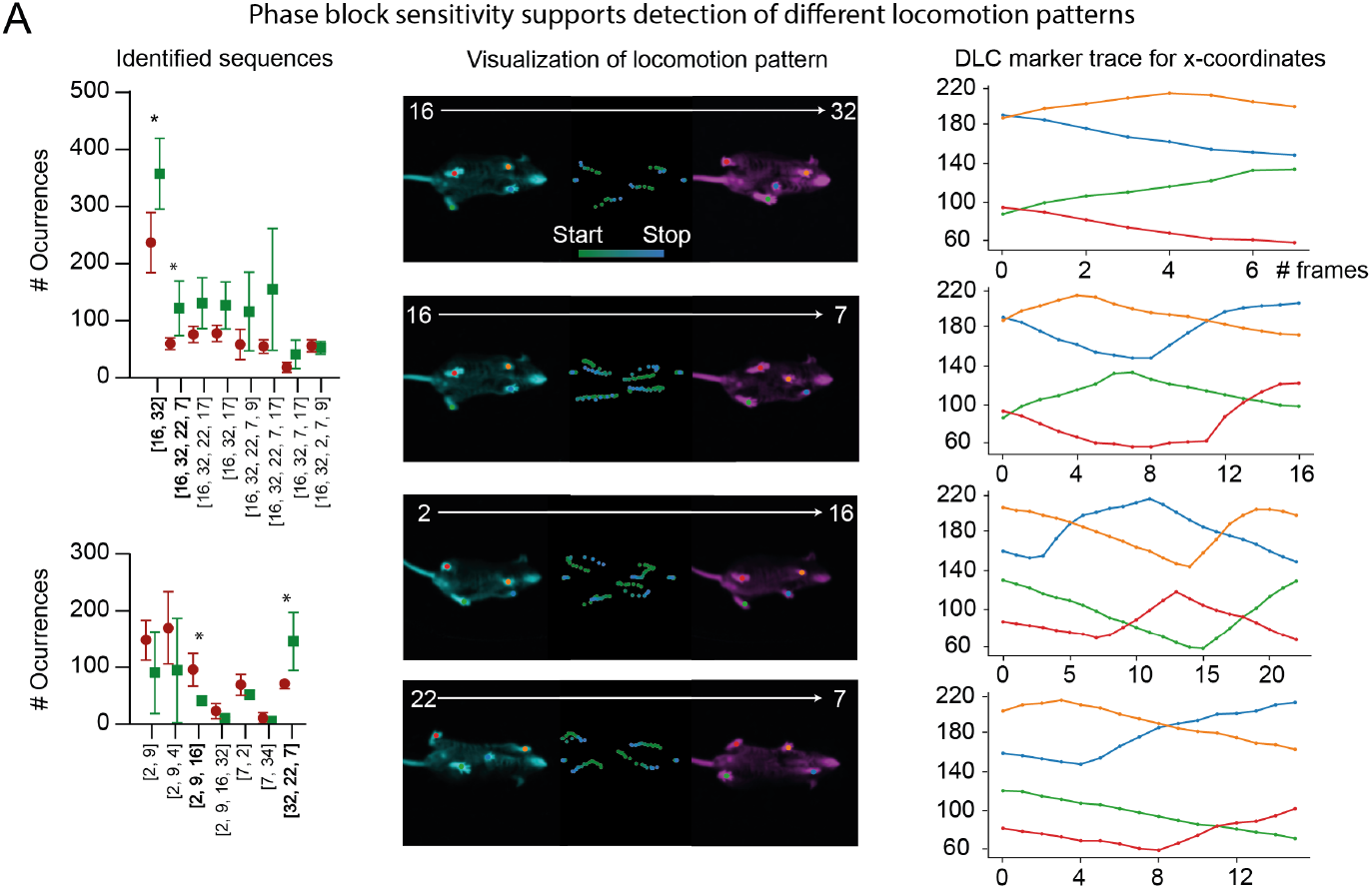
(A) Identfication of the locomotion structure in both groups (left) and visualization of four significant different patterns (middle, right).

Given the results shown in Figure 3 C, we used the representation learned by VAME to identify locomotion patterns within the *Walking* community. Briefly, we built an algorithm that first identified reoccurring motif patterns in the community and then counts their occurrences. We detected 22 sub-patterns, which were used by all animals. Within these sub-patterns, we were able to identify four specific sub-second sequences that were significantly more used in either wt or tg mice (unpaired t-Test). The strongest sub-pattern for wt animals is the sequence [16, 32] and [32, 22, 7], which can be also seen in the difference graph as a strong transition. For the tg, the strongest pattern involves the sequence [2, 9], which again can be seen in the difference graph as a strong transition for the tg group. These results demonstrate the capabaility of VAME to robustly capture patterns for transitions between motifs, especially for locomotion behavior based on its phase sensitivity.

### 5.10 Community transitions

**Figure S.8:**
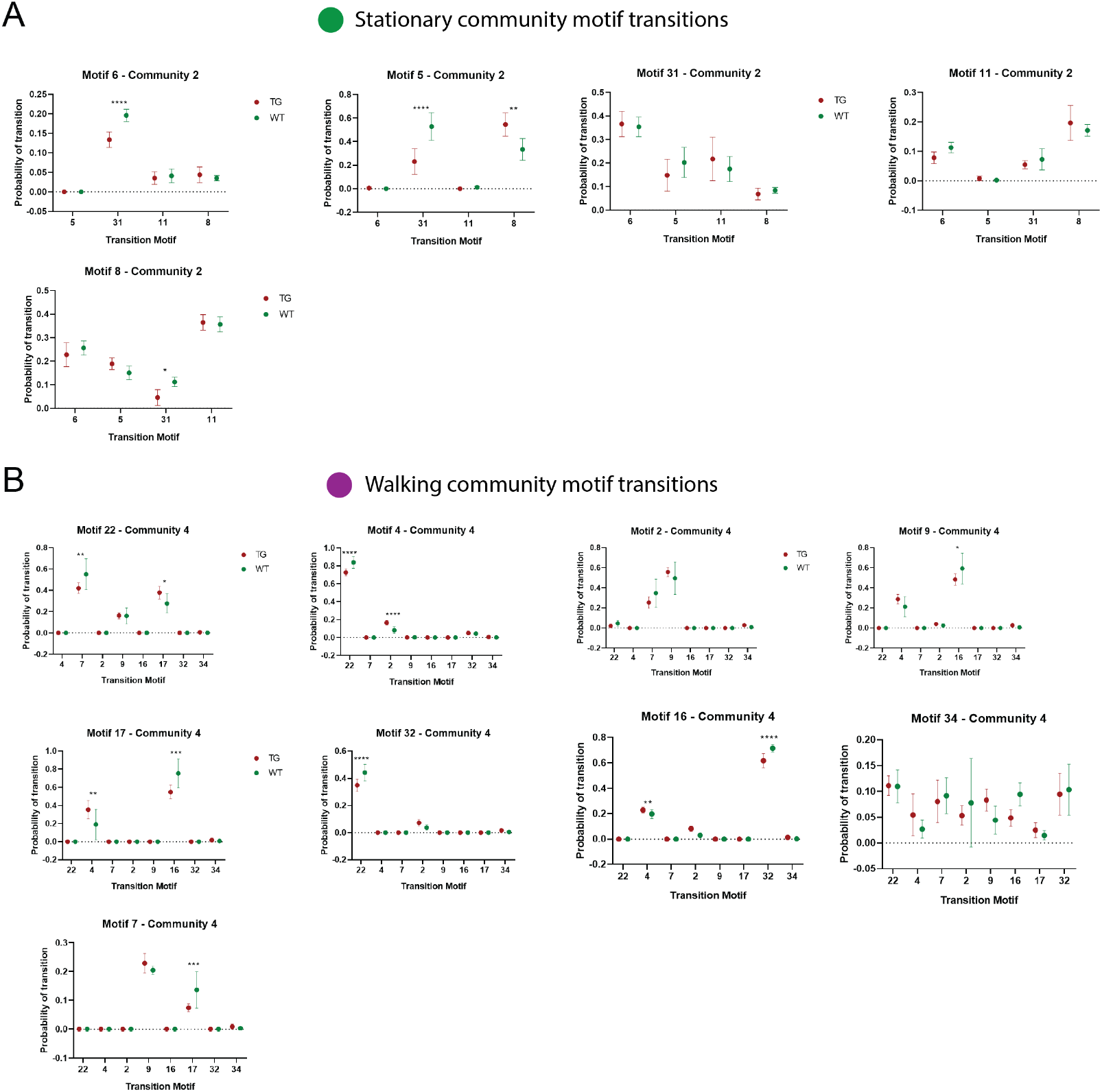
Transitions of motif within the *Stationary* and *Walking* community.

**Figure S.9:**
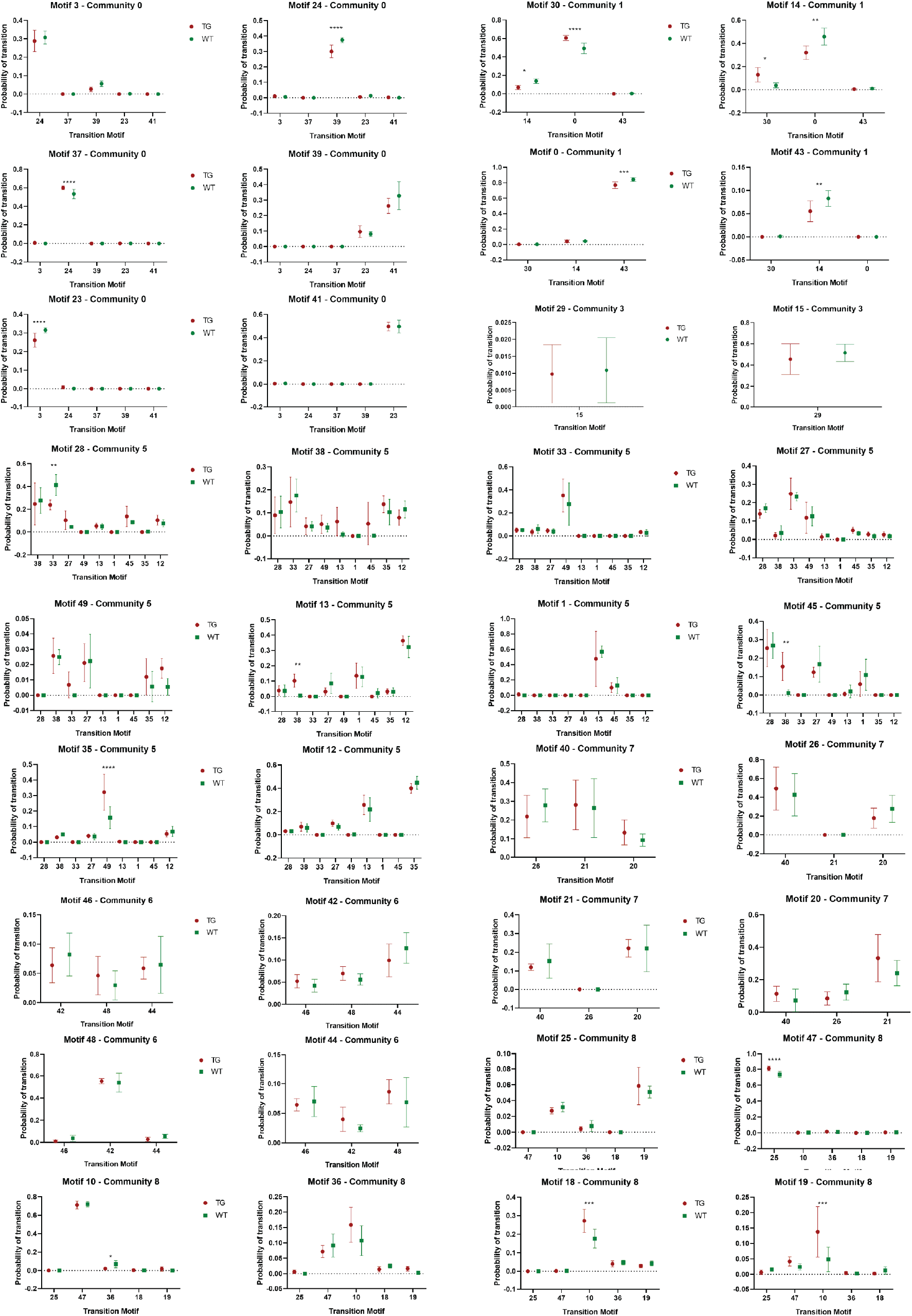
Within community transition statistics.

